# CropQuant: An automated and scalable field phenotyping platform for crop monitoring and trait measurements to facilitate breeding and digital agriculture

**DOI:** 10.1101/161547

**Authors:** Ji Zhou, Daniel Reynolds, Thomas Le Cornu, Danny Websdale, Simon Orford, Clare Lister, Oscar Gonzalez-Navarro, Stephen Laycock, Graham Finlayson, Tim Stitt, Matthew D. Clark, Michael W. Bevan, Simon Griffiths

## Abstract

Automated phenotyping technologies are capable of providing continuous and precise measurements of traits that are key to today’s crop research, breeding and agronomic practices. In additional to monitoring developmental changes, high-frequency and high-precision phenotypic analysis can enable both accurate delineation of the genotype-to-phenotype pathway and the identification of genetic variation influencing environmental adaptation and yield potential. Here, we present an automated and scalable field phenotyping platform called CropQuant, designed for easy and cost-effective deployment in different environments. To manage infield experiments and crop-climate data collection, we have also developed a web-based control system called CropMonitor to provide a unified graphical user interface (GUI) to enable realtime interactions between users and their experiments. Furthermore, we established a high-throughput trait analysis pipeline for phenotypic analyses so that lightweight machine-learning modelling can be executed on CropQuant workstations to study the dynamic interactions between genotypes (G), phenotypes (P), and environmental factors (E). We have used these technologies since 2015 and reported results generated in 2015 and 2016 field experiments, including developmental profiles of five wheat genotypes, performance-related traits analyses, and new biological insights emerged from the application of the CropQuant platform.

## Introduction

The great American wheat breeder and agricultural innovator Orville Vogel once stated, the plant we are looking for is in our plots, but we have to be there when it is. In order to select varieties with greater yield potential and enhanced environmental adaptation, agricultural practitioners, including breeders, farmers and crop researchers, have been optimising trait combination since the beginning of agriculture^1,2^. Four decades after temporary success in ensuring global food security^3^, we are now facing an even bigger challenge to feed generations to come^4^. Due to a narrowing range of available genetic diversity of modern crop germplasm collections^5^ and increased fluctuations in growing conditions^6^, there is increasing emphasis placed on exploiting new sources of genetic variation to enhance environmental adaptation and sustainable yield in crop landraces and wild relatives^7^. To identify and assess these types of traits, multiple regular measures of crop growth and development are required to quantify subtle and dynamic phenotypes from many plots in different growing environments, demanding new screening technologies to integrate field environmental datasets with multi-scale phenotypic analyses to understand genotype-by-environment interactions (GxE) and associate them to genetic variation^8–10^.

In contrast to current field phenotyping methodologies, which are still involving laborious manual scoring and relatively subjective selection, modern genetic and genomics techniques are being rapidly deployed in breeding and crop research to identify and utilise traits such as improved stress tolerance and disease resistance^11^. For example, quantitative trait locus (QTL) analysis and genome-wide association studies (GWAS) are used to identify loci^12^, whole genome sequencing used to reveal gene content and genetic variation^13^, and marker-assisted selection (MAS) and genomic selection used to breed new lines with favourable alleles^14,15^. Therefore, field phenotyping and the integration of environmental data with meaningful phenotypic analyses are a key bottleneck that limits the potential of recent advances in crop genetic and genomic technologies^16,17^.

Remote sensing platforms^18^ and open image-based analytics software libraries^19^ start to enable researchers, breeders and agronomists to develop new approaches to understand and improve crop performance. For instance, unmanned aerial vehicles (UAVs) and light aircraft are being used to study crop performance and field variability^20,21^. Satellite imaging^22–24^ and ground-based portable devices^25,26^ have been applied to take snapshots of crop growth to estimate yield-related traits using canopy photosynthesis rate and normalised difference vegetation indices (NDVI). Field-based agricultural vehicles have been developed to capture physiological and developmental traits during the growing season^18,27^. Finally, large imaging platforms equipped with 3D laser scanners (e.g. near-infrared laser lines and Light Detection and Ranging, LiDAR) and multi– or hyper-spectral sensors are applied to automate crop monitoring of a fixed number of pots or plots, either in the field^28,29^ or in greenhouses^12,30^. Although these advances are making important contributions to the research domain, there are limitations and challenges associated with their usage such as high costs, restricted mobility and scalability, limited frequency of screening, and inadequate software tools for phenotypic analyses^31,32^. In particular, while satellite imagery and UAVs are capable of screening tens of thousands of plots at multiple locations, their applications are subject to civil aviation rules, low spatial resolution and bad weather conditions such as heavy rainfall, strong wind and cloud coverage. Ground-based portable devices and vehicles have shown greater mobility and high-resolution field data between multiple sites; however, they require experienced specialists to operate, limiting their applications to infrequent phenotyping. Still, stationary large imaging platforms are providing key data on dynamic crop growth and GxE interactions; but their scale of operation is restricted and they are relatively expensive for less well-funded research laboratories to access. Furthermore, they mostly rely on proprietary analytics software for data management and trait analysis, requiring ongoing licensing maintenance to use software products and extra fees if tailored functions are needed. For these reasons, it is challenging for researchers and breeding communities to adopt new phenotyping approaches due to their expenses, lack of suitable software, limited scope of operation and maintenance costs^33^. The ability to facilitate crop improvement programmes at multiple scales and locations is still limited.

To enable the next-generation breeding and associated crop research^34^, affordable field phenotyping technologies need to be developed. New methods should exploit up-to-date remote sensing technologies together with state-of-the-art computer vision and software solutions, to equip researchers with diverse tools for multi-scale field phenotyping needs. The work described here aims to address these challenges by the development of an automated field phenotyping platform called **CropQuant**, which integrates cost-effective hardware with open source software capable of complex analytic solutions. We demonstrate applications of CropQuant (CQ) through multiple growth phenotypes measurements based on defined wheat genotypes over two growing seasons. Dynamic predictive growth models were also created to forecast the performance of wheat genotypes under varied environment conditions.

## Materials and Methods

### An open and low-cost hardware design

In order to carry out automated phenotyping, we have deployed many low-cost crop monitoring CQ workstations *(terminals)* operating jointly via a preinstalled or self-operating infield wireless network. **Figure 1** shows the system architecture of the CQ platform. A terminal is centred around a simple single-board computer (**Fig. 1a**), running a customised Python-based analytic software package on the Linux *Debian* operating system (OS), integrating *Pi* or USB camera sensors (e.g. red-green-blue (RGB) or no infrared filter, NoIR), climate sensors (ambient and soil-based), sensor circuit boards, with wired or wireless data communications (**Figs 1b&c**). The design of the CQ platform is driven by the concept of Internet of Things (IoT)^35,36^ as well as how to utilise hardware and software resources that are widely available, so that crop phenotyping solution can be scalable and affordable for the communities.

**Figure 1.**
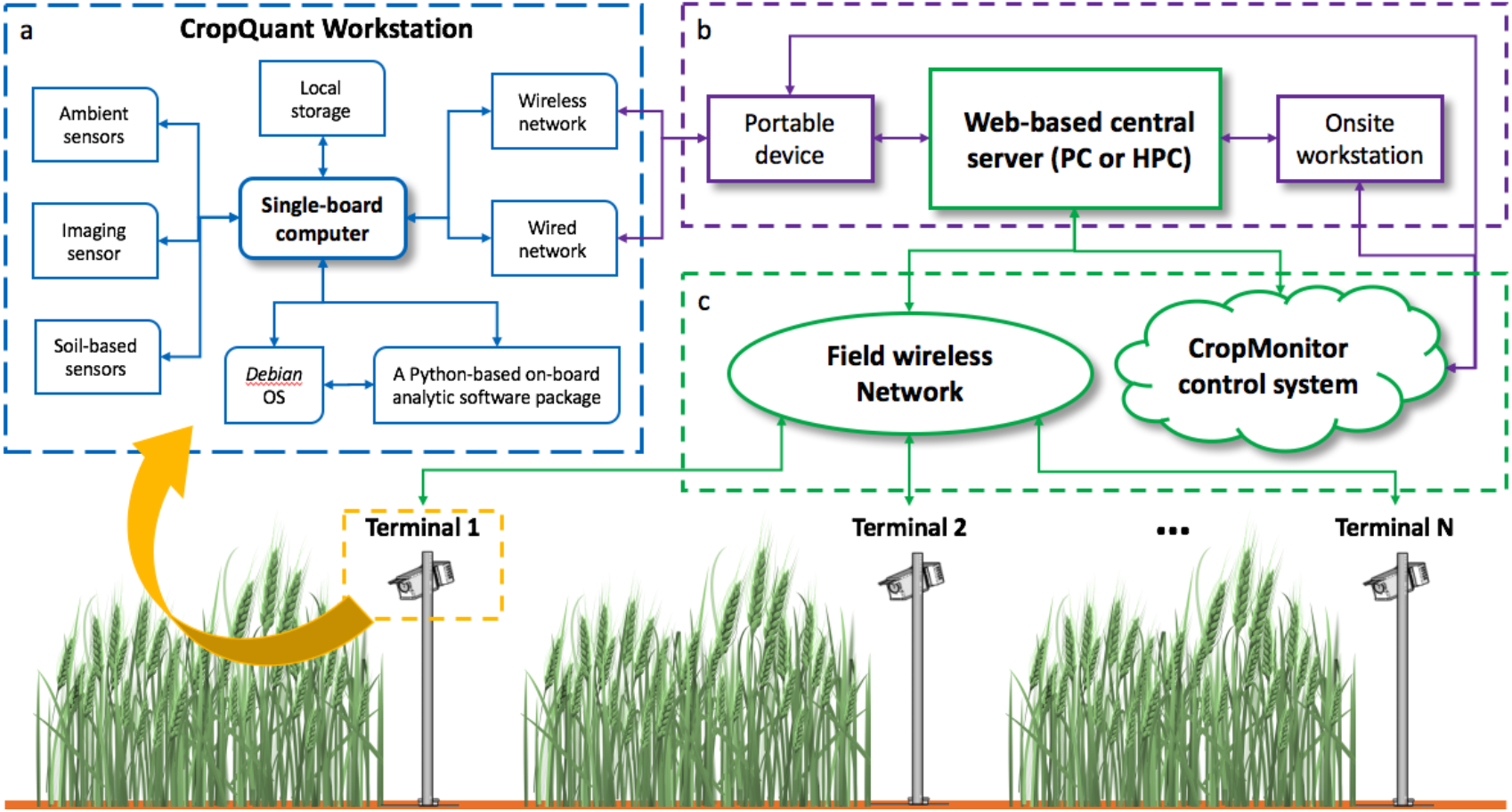
A high-level system architecture of the CropQuant platform. **(a)** The hardware and software design for a CQ workstation, including a single-board computer, climate sensors, a tailored circuit board to integrate sensors, an imaging sensor, local USB storage, wired and wireless network components, *Debian* operating system, and custom-made Python software package for crop images and climate data collection. **(b)** Data communications between CQs and either a portable device in the field or an onsite PC workstation. **(c)** The network setting that integrates CQ terminals, infield wireless network, and the CropMonitor control system.

We have used these technologies in field experiments since 2015. **Figure 2** shows an experimental scale of how CQ workstations were deployed for onsite and offsite wheat assessment experiments in 2015 and 2016. To keep costs low so that the technology can be adopted by the communities easily, we have developed different CQ versions (**Figs 2a-d**). For example, an all-in-one CQ (**Fig. 2a**) uses a *Raspberry Pi* 2 single-board computer to control internal hardware (Figs 2b), including (1) a *Pi* camera sensor for time-lapse crop photography, (2) a tailored circuit board (**Fig. 2c**) to integrate climate sensors for collecting environmental data (i.e. soil temperature and moisture, ambient temperature and humidity, and light levels), (3) a USB WiFi dongle (or a radio transmitter) for data transfer and remote interactions, and (4) a USB flash drive for local data storage. **Supplementary Figure 1** demonstrates how an all-in-one CQ was used in field experiments.

**Figure 2.**
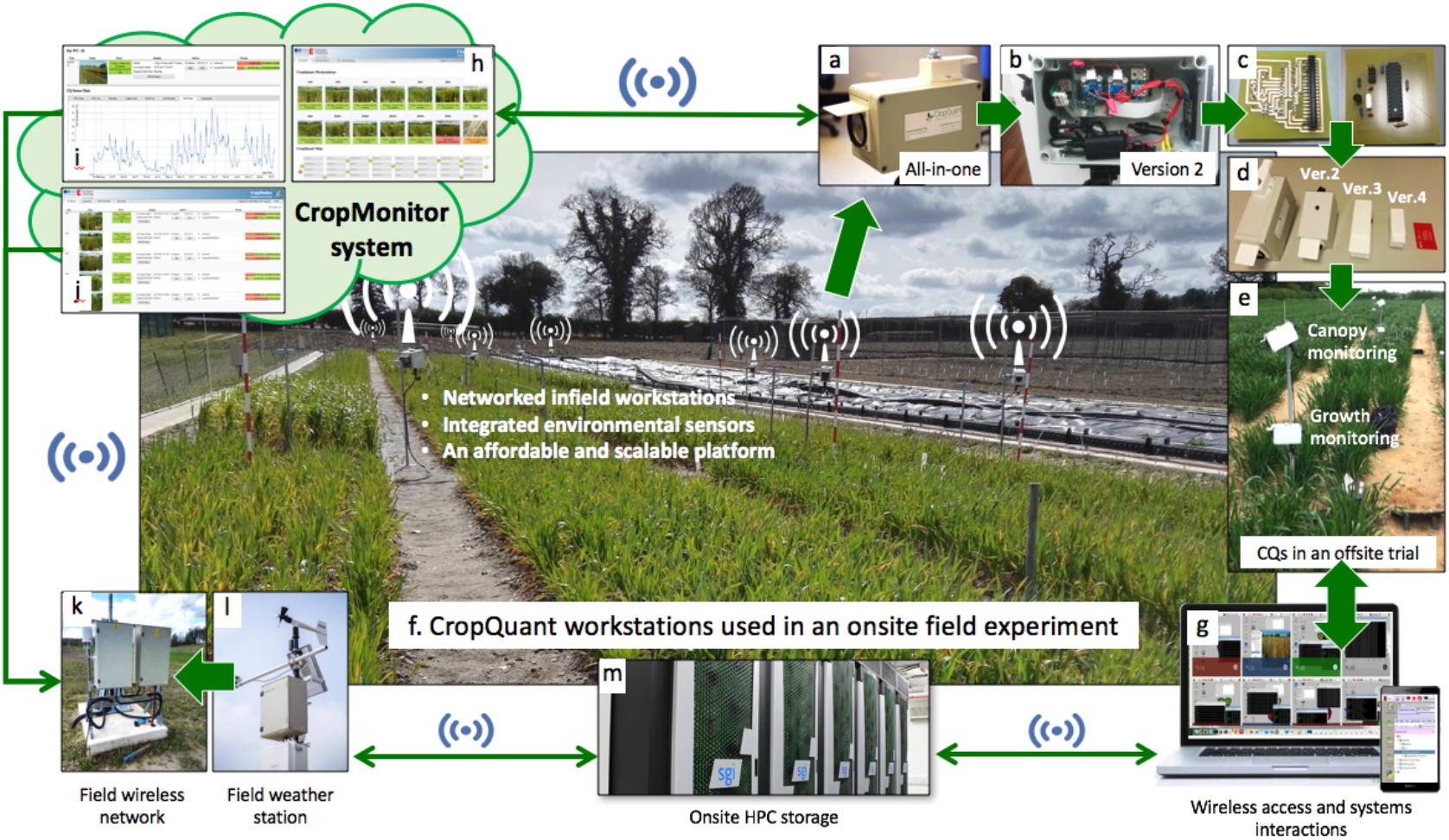
The CropQuant phenotyping platform used in field experiments. **(a)** An all-in-one CQ workstation. **(b)** The internal hardware design of a version 2 CQ workstation. **(c)** A custom-made circuit board that integrates low-cost climate sensors. **(d)** Different versions of CQ built for dissimilar monitoring tasks. **(e)** CQs used in offsite field experiments, powered by batteries and solar panels. **(f)** The CQ platform established for onsite field experiments, powered by 5V/2A power supplies. **(g)** Realtime crop inspection using either a portal device in the field or a PC in an office. (h-j) the CropMonitor control system administers CQ terminals and records information such as online or offline status, operational mode, daily representative crop image, micro-environment for the plot region, and computational resource for each connected CQ terminal. **(k)** An infield WIFI system installed for field trials. **(l)** A commercial field weather station. **(m)** HPC clusters used for durable data storage and trait analysis.

Besides the relatively costly all-in-one version ($230-$240 to build), other more specialised versions (**Fig. 2d**) were much cheaper to produce. For example, a business card size CQ (version 4, $80-90) uses a *Pi* Zero computer and is tailored for crop photography. A version 2 CQ (**Fig. 2b**, $170-180) can mount different sensor groups (ambient or soil-based) for assessing agronomic characterisation and crop adaptation. With hardware modularity in mind, we have tested a range of single-board computers (e.g. the *Raspberry Pi* series, Intel’s Galileo and Edison) for performing simple infield image analysis as well as integrating modular components. Although we finally chose *Pi* computers due to its performance-to-price ratio and extensive community support, the platform can operate on other single-board computers, if *Pi* computers are not available. For the peripheral hardware, we used off-the-shelf weatherproof containers (IP67 rating) together with micro-USB and Ethernet couplers (IP66 rating) to ensure environmental endurance and outdoor deployment. **Supplementary Figure 2** shows the hardware design of a version 2 CQ. A full hardware list and a construction manual are included in **Supplementary Note 1**.

### Offsite self-operating mode

**Figure 2e** demonstrates an offsite field experiment using the CQ platform in 2015, where 10 field workstations were deployed to monitor canopy development (**Movie 1**) and crop growth (**Movie 2**) on one-metre wheat plots. CQ devices were powered by lead acid batteries with trickle charging from solar panels. To operate the device with minimal energy requirements, we have implemented a headless access mode to carry out wired data transfer. Besides the programmed imaging task, the system was only wakened if an Ethernet connection (i.e. local area network, LAN) was established. Offsite CQs were self-operating and used to perform image-based phenotyping. The infield imaging script running on CQs can be seen in **Supplementary Note 2**.

### Onsite networking mode

For onsite experiments (**Fig. 2f**), CQs were powered by 5V/2A power supplies and connected to a field WiFi network. There were 14 networked CQ terminals (21 at peak time, with two dedicated for tiller abortion studies) jointly operating in 2016, monitoring 12 six-metre wheat plots to study performance-related traits and yield production. Although not thoroughly tested, for pre-installed WiFi network, the scale of the CQ platform can be increased by adding more standard routers to allow more CQ connections. We have added new functions to networked CQs including wireless control, programmable imaging, and on-board quality control (Online Methods). For instance, end-users can access CQ workstations remotely for real-time inspection, either using a portable device (e.g. a tablet or a smartphone) in the field or an onsite office computer (**Fig. 2g** and **Supplementary Fig. 3**). They can check the field in different regions to: (1) review historical crop images, (2) initiate new experiments, and (3) transfer crop-climate data to external computing storage.

**Figure 3.**
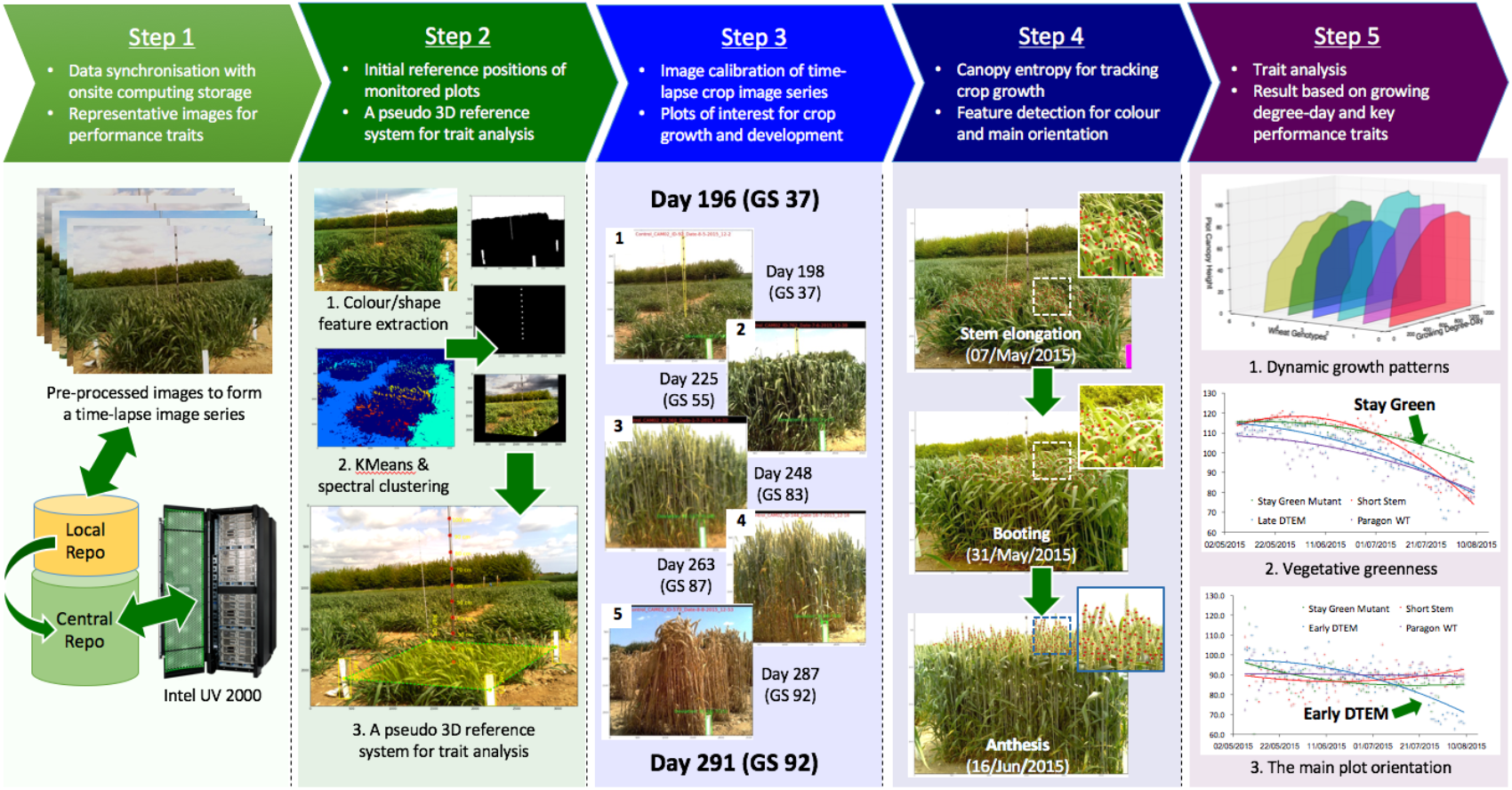
The high-throughput analysis pipeline established for batch processing and measuring performance-related traits. **(1)** *Step 1,* high-quality crop images were selected by the selection algorithm and stored in both local and central repositories. **(2)** *Step 2*, initial reference positions of monitored plots were detected by the referencing algorithm, which also calculated the pixel-metric conversion rate. **(3)** *Steps 3* and 4, the performance-related trait analysis algorithm was designed to track plots of interest and conduct trait analyses to quantify the canopy region and performance-related traits. (4) *Step 5,* traits such as dynamic crop growth patterns in relation to thermal time (degree-day), vegetative greenness (0255), and the main plot orientation (0^o^-180^o^) were quantified and illustrated.

These monitoring activities are administered by our web-based control system, CropMonitor (**Figs 2h-j** and **Supplementary Fig. 4**), where the status of each CQ terminal is updated constantly with information such as online or offline status, operational mode (e.g. green for operating, amber for idle, and red for operation error or ending tasks), representative daily images, micro-environment readings, and the usage of computing resources (i.e. CPU and memory). Furthermore, the CropMonitor system can support a range of tasks. For example, when deploying CQs in the field, CropMonitor can activate live streaming between a CQ terminal and a smart device (e.g. a smartphone or a tablet) to enable the calibration and installation of CQ devices (**Supplementary Fig. 5**). During the experiment, the CropMonitor system can establish a mesh network (based on all-in-one CQs) to support data communications between the field and external servers (Online Methods and **Supplementary Note 3**). Users can reposition CQ terminals at any time to change or initiate new monitoring tasks. Notably, the IoT-style setting can improve the mobility of the CQ platform. For example, **Supplementary Figure 6** shows a speed breeding experiment^37^ monitored by CQs over a 75-day period in 2017, which was accomplished by moving CQs to indoor, i.e. a growth chamber (the speed breeding condition) and a glasshouse (the control condition).

**Figure 4.**
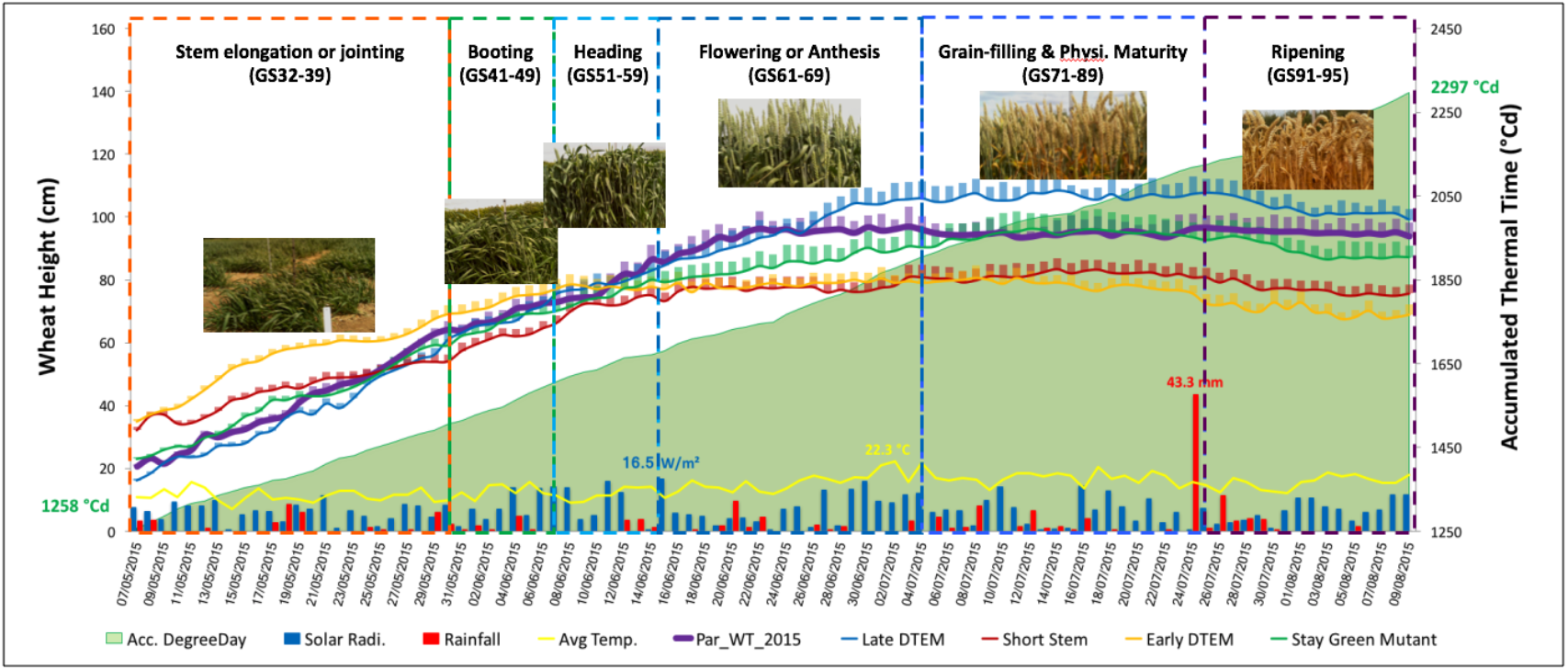
The performance of five wheat NILs monitored by the CQ platform to identify dynamic developmental profiles. Five wheat NILs (Late-DTEM, Early-DTEM; Short, Stay-Green, and *Paragon* WT) and their dynamic performance in relation to environmental factors such as solar radiation, rainfall, and temperature, during the 95-day monitoring period. Six growth stages of *Paragon* WT were used as reference. Accumulated thermal time in degree-day units was computed for comparison.

**Figure 5.**
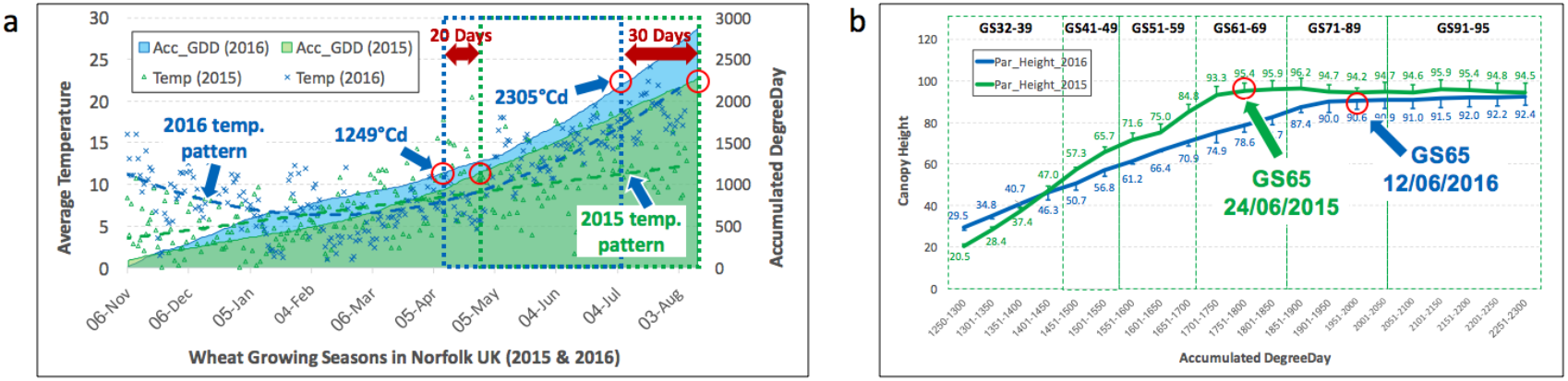
Recognising subtle and dynamic developmental variances for wheat genotypes under different climate patterns using the CQ platform. **(a)** Different temperatures and accumulated degree-day patterns recorded in 2015 and 2016. **(b)** Aligning and comparing growth curves of *Paragon* WT in 2015 and 2016 within similar growth stages and the degree-day period (1250-2300 °Cd).

**Figure 6.**
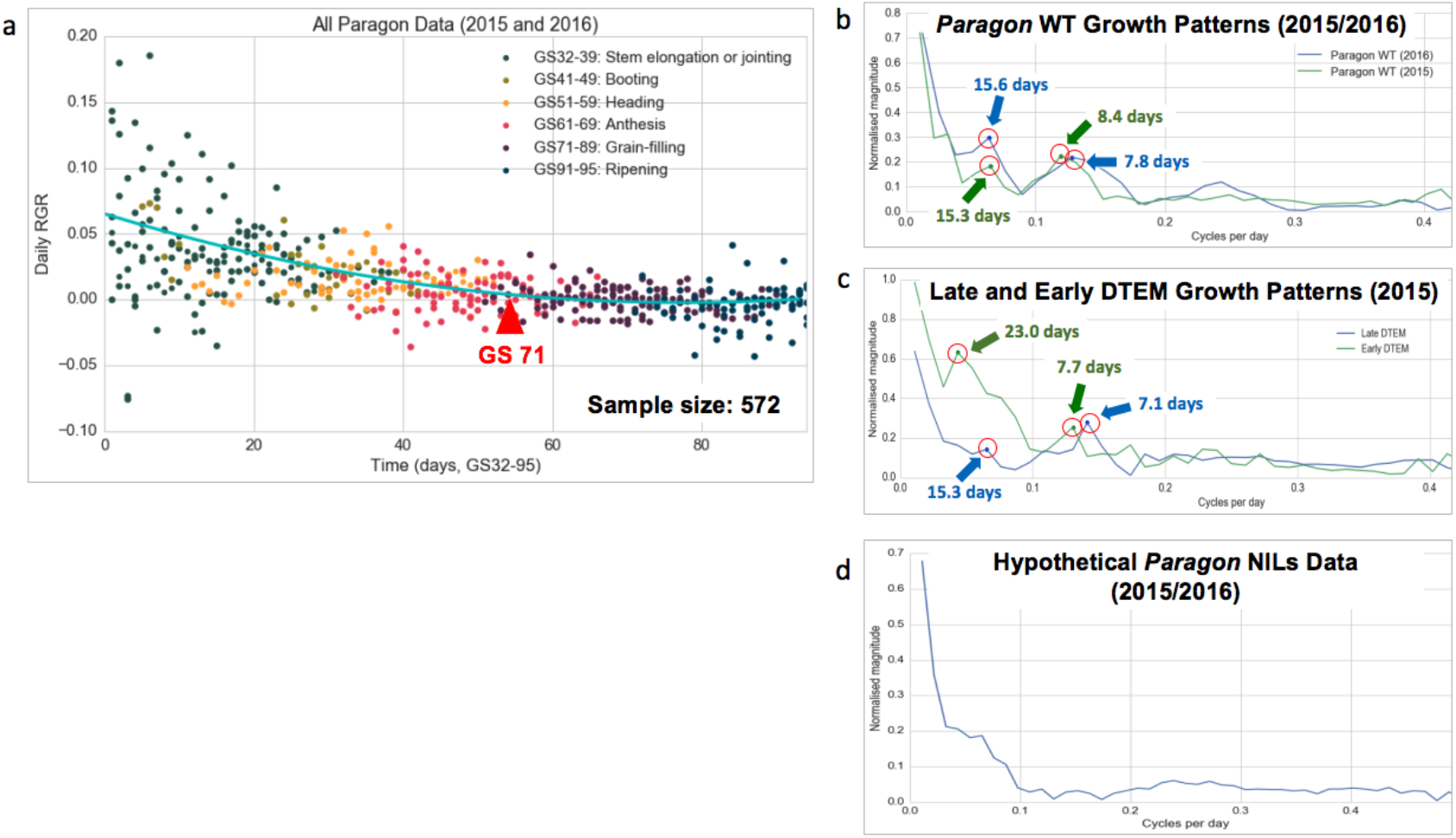
Extracting underlying growth patterns from continuous phenotypic data using a fast Fourier transform (FFT) **(a)** RGR (growth % of the previous day) is used to present the daily growth rate of all the NILs at different growth stages. **(b)** The underlying growth patterns for *Paragon* WT in 2015 and 2016 after the FFT conversion. **(c)** The underlying growth patterns for Late-DTEM and Early-DTEM, after the FFT conversion. **(d)** Using a hypothetical *Paragon* NILs growth data (merging all the NILs) to study the growth pattern.

To increase the scalability of the phenotyping platform, we recently developed a mesh network system to connect infield terminals with or without any pre-installed network infrastructure. **Supplementary Figure 7** illustrates the network topology, where all-in-one CQs are operating jointly as backbone *routers* (i.e. cluster servers), running both dynamic host configuration protocol (DHCP) and virtual network computing (VNC) servers for networking. Cheaper CQs (versions 2-4) are connected to the *routers* as terminals (i.e. *end devices*). Depending on the number of *routers* and the coverage of WiFi dongles (25-30 metres) or radio transmitters (several hundred metres), the field mesh network can be expanded or downsized flexibly. We dedicate an all-in-one CQ as a *coordinator* to control data communications between terminals and external networks. Meanwhile, computing tasks such as image selection, quality control and initial data annotation are distributed to terminals to reduce computational burden of the in-depth trait analysis (**Supplementary Note 4**).

**Figure 7.**
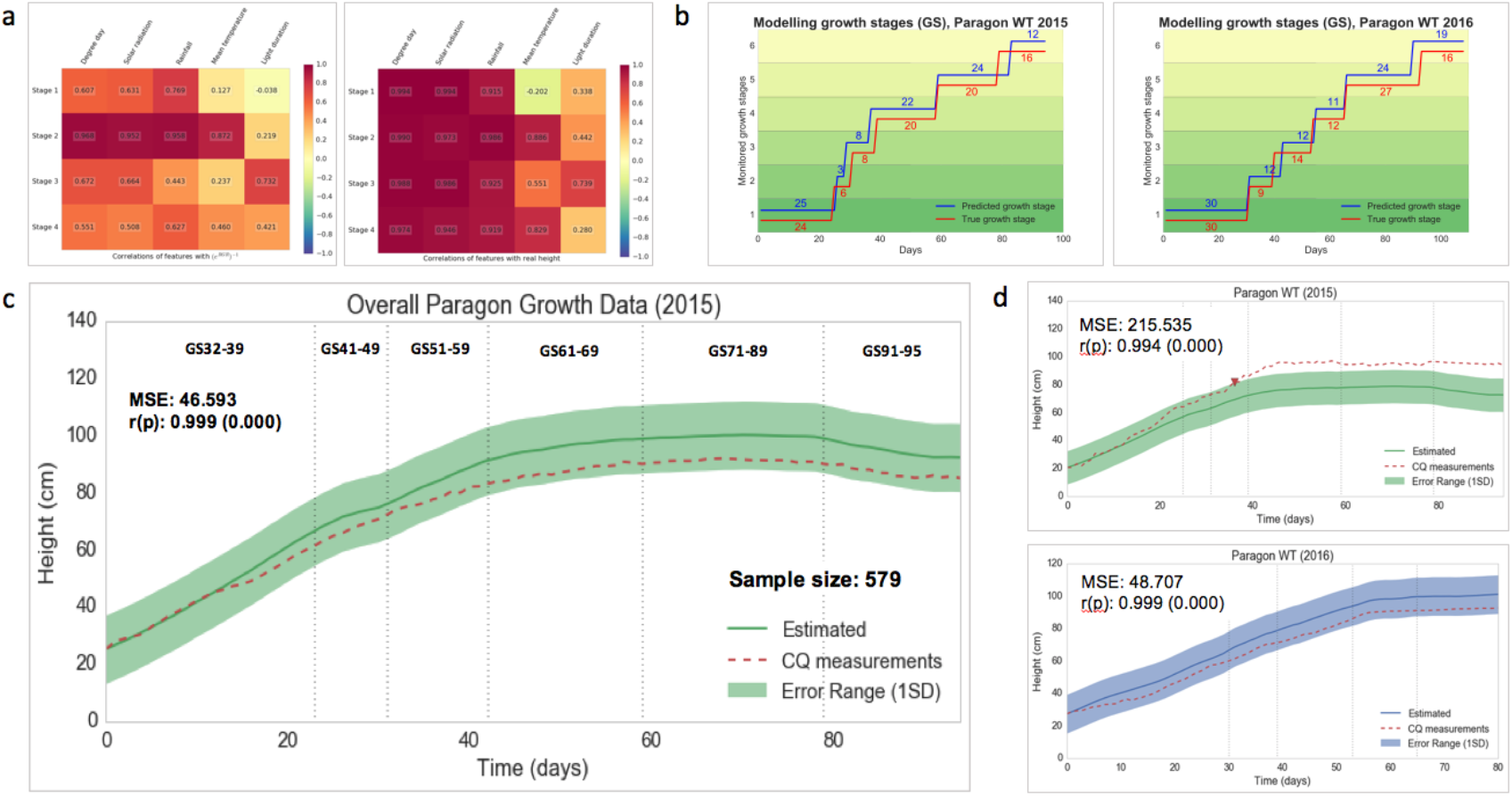
Establishing a GxE predictive model to forecast the performance of wheat genotypes under different climate patterns. **(a)** Applying a correlation model to identify highly correlated environmental factors, including thermal time, solar radiation, rainfall and growth stage duration using *Pearson* correlation (p<0.01). **(b)** A growth stage-based predictive model applied to estimate the key growth stages of *Paragon* WT in 2015 and 2016 compared with manual scoring. **(c)** A global growth model comparing real growth curve measured by CQ (red dotted line) with estimated growth curve (green line). **(d)** Warning messages triggering mechanism to alert users if crop growth is outside the safe bounds (±1SD) of the estimated growth region.

Additionally, to verify the outputs of low-cost remote sensors integrated in CQs, we have utilised the meteorological outputs of a commercial weather station (**Figs 2k&l**), including temperature, rainfall, photosynthetically active solar radiation, wind speed and relative humidity. Phenotypic and climate datasets were saved on an onsite high-performance computing (HPC) cluster (SGI UV2000 system with Intel Xeon cores) for durable data storage (**Fig. 2m**). Since the application of the CQ technology, we have successfully accomplished three tasks essential for the next-generation field phenotyping^17^: (1) *continuous monitoring* via time-lapse crop photography, (2) *infield evaluation* through networked terminals and the CropMonitor system, and (3) *efficient data transfer* using distributed computing and wireless data communications through an infield network (**Supplementary Fig. 8**).

### The high-throughput analysis pipeline

In order to enable accurate delineation of the genotype-to-phenotype pathway and identify genetic variation influencing environmental adaptation and yield potential, we chose a high-frequency (two-three times per hour) and high-precision (2592x1944 pixels per image) phenotyping approach to monitor the morphological change of crops. After the crop photography phase, we exploited open image processing and machine learning libraries such as OpenCV^38^, Scikit-learn^39^ and Scikit-image^40^ and developed an automated analysis pipeline to extract biologically relevant outputs.

We designed the analysis pipeline to be executable on either a workstation PC or an HPC cluster. Firstly, to arrange the collected image series, we have developed a selection algorithm to choose representative images based on their size, clarity^41^, imaging dates and genotypes (Fig. 3, *Step* 1). Only high-quality images were retained for trait analysis (Online Methods and **Supplementary Note 5**). All datasets were archived in a central repository such as HPC clusters for future reference. Then, we developed a referencing algorithm to define the location of a monitored plot over time (Fig. 3, *Step* 2). In real-world agricultural and breeding situations for which CQs are deployed, strong wind, heavy rainfall, irrigation and chemical spraying can lead to modest camera movements, causing cross-reference problems when comparing trait analyses for a plot over time (**Movie 3**). To resolve this issue, we have designed the referencing algorithm to identify the initial plot location so that each image in the series can be transferred to the same position for comparison. For example, the algorithm detects 2D coordinates of white reference canes installed in the plot and dark markers on a ranging pole for height scales using colour- and shape-based feature selection^39^. Then, it classifies pixels into five groups to represent the canopy space and background objects such as wheel tracks, sky, and the reference canes using simple k-means^42^ and spectral clustering^43^ algorithms. Finally, a pseudo 3D reference system is established to record important coordinates of the plot region, the canopy space, and height markers, together with converting measurements from pixels to metric units such as centimetres (Online Methods and **Supplementary Note 6**).

Following Step 2, we integrated the initial reference location into a performance-related trait analysis algorithm. For a given image series, the algorithm applies an adaptive intensity and gamma equalisation method^44^ to minimise colour distortion caused by varied field illumination. Then, it tracks geometric differences^38^ between the monitored plot and the reference location. If the plot location has changed in a given image, a geometric transformation method^45^ will be applied to reposition the image, removing areas outside the plot region and may or may not generate a black bar to the top of the image (Fig. 3, *Step* 3). Within the plot region, the algorithm detects the visible part of the ranging pole (**Movie 4**) as well as the canopy space for height measurement. For instance, to measure the height of the canopy, an entropy-based texture analysis is used to determine whether the canopy region is changing between two consecutive images using gray-level co-occurrence matrices (GLCM)^46^. If positional changes (moving up or down, depending on growth stages) are identified, the canopy height is recorded and corner-featured points^47^ are detected (Fig. 3, *Step* 4), generating many red pseudo points casting in the canopy region for measuring canopy height (Movie 5 and **Supplementary Note 7**). These pseudo points can also be used to represent the tips of erect leaves at stem elongation or jointing (the Zadoks scale^48^, growth stages, GS 32-39), reflective surfaces of curving leaves or crop heads between booting and anthesis (GS 41-69), and corner points on spikelets during senescence (GS 71-95). Using the trait analysis algorithm, we have computed the dynamic height changes to present growth patterns for different wheat genotypes (Fig. 3, *Step 5.1*).

### Developmental related trait measurements

In addition to the canopy height, we also developed functions to calculate other traits. For example, vegetative greenness is calculated based on the normalised greenness value (0-255) for a given plot over time. The output was used to assess the change of green biomass and vegetation period. We used this trait to evaluate a Stay-Green mutant (prolonged green leaf area duration with delayed leaf senescence; Fig. 3, *Step 5.2*). The main orientation of a plot (0-180^o^) is also quantified based on edge detection methods^49^, representing the alignment of stems to estimate the change of stem rigidity (Fig. 3, *Step 5.3*). Using this trait, we have identified lines with higher lodging risk either during ripening or when interacting with heavy rainfall or strong wind (Online Methods and **Supplementary Table 1**).

## Results

### Use case 1 – Monitoring five wheat NILs

The diverse environments for which wheat has been adapted to grow provide opportunities for us to explore the dynamic interactions between genetic diversity and phenotypic traits under varied environmental conditions^50^. To test the CQ platform, we chose wheat near-isogenic lines (NILs, Online Methods) to examine a number of key performance-related phenotypes in the same *Paragon* (a UK spring wheat variety) genetic background^51^. **Figure 4** demonstrates five dynamic developmental profiles generated from the experiment between May and August 2015, a 95-day period. The experiment was conducted in plots in a field which is 2.1 miles away from Norwich Research Park UK (see the plot layout in **Supplementary Table 2**) and all five NILs were monitored twice per hour. The genotypes were: (1) Late-DTEM (days to ear emergence^48^, the number of days between sowing and ear emergence; *late* means GS55 is delayed), with *Ppd-1* loss of function (lof); (2) Early-DTEM (GS55 is moved forward), with *Ppd-D1a* photoperiod insensitivity; (3) Short stems, *Rht-D1b* semi dwarfing; (4) Stay-Green, a stay green mutant; and (5) *Paragon* wild type (WT).

To compare the performance of the NILs, we used *Paragon* WT as the reference line and highlighted six key growth stages, from stem elongation or jointing (GS 32-39) to ripening (GS 91-95). The thermal time (degree-day, ^o^Cd, using a 0°C base^52^) was also used as a heuristic tool^53^ to normalise the crop growth. The five growth curves (1258-2297 °Cd) approximately followed a sigmoid curve. At the beginning of the experiment, *Ppd-D1a* NIL (Early-DTEM, coloured amber) was already at the end of the jointing stage (GS37-39) and hence was the first to reach a maximum height; whereas *Ppd-1* lof (Late-DTEM, coloured blue) was the last to increase in height. By cross-referencing developmental profiles based on six growth stages, we noticed that although *Ppd-D1a* and *Rht-D1b* (Short-Stem, coloured red) had similar maximum heights (83.4cm and 80.6cm), the latter displayed a relatively steady rate of increase in stature. *Ppd-1* lof’s growth was the most delayed line, resulting in an extended period of vegetation, stem extension, and overall time to ear emergence. As this genotype has received the most thermal time units, it was the tallest line in the field experiment. Although all NILs experienced some degree of height reduction due to a significant storm on 24^th^ July 2015, *Paragon* WT (coloured purple) presented a much lower lodging risk, as it maintained its height afterwards. To verify the phenotypic observation, we scored heading dates and canopy heights manually on the same plots and obtained a Pearson correlation coefficient of 0.986 (**Supplementary Table 3**).

We summarised different temperatures and accumulated degree-days (ADDs) in both 2015 and 2016 growing seasons (**Fig. 5a**). As the average temperature in 2015 is much lower than in 2016, we used a fixed ADD period (1250-2300 °Cd) to segment crop growth under different climates. Within the same ADD period, Figure 5b shows dissimilar growth curves of *Paragon* WT (**Supplementary Table 4**). The 2015 curve was much steeper during stem elongation (GS32-59, 1250-1750 °Cd), possibly reflecting the cold spring. Flowering half complete (GS65) was reached on 24^th^ June 2015 (an early drilling late maturity mode). While the 2016 curve (values are the means of two biological replicates) had a steadier and extended development due to a warm spring. GS65 was reached on 12^th^ June 2016, 12 days ahead of 2015 (a late drilling early maturity mode). Using the CQ platform, not only have we collected high-frequency crop climate datasets, we also could identify dynamic developmental variances for genotypes under different climate patterns.

### Use case 2 – New biological insights into growth patterns

High-frequency and high-resolution deep phenotyping has already been employed in human disease research to reveal the underling mechanisms of individual’s disease^54^. In plant research, the similar approach is being adopted to characterise phenotypes of plant responses to environmental challenges for field experiments^55^. While applying CQs in wheat assessment experiments, we have explored new biological insights into dynamic growth patterns using the large phenotypic data captured in the field.

**Figure 6** presents some preliminary results of how we utilised the high-frequency phenotypic data to extract underlying growth patterns for *Paragon* genotypes. Initially, we calculated daily relative growth rates (RGR, comparing with the previous day) for the five NILs monitored in 2015 and compared them with two *Paragon* WT measured in 2016. All RGR data were aligned by the associated growth stages for comparison, showing that all lines were active from jointing to flowering and became inactive after grain-filling (**Fig. 6a**). After that, to study the change of RGR during the growth stages (GS32-69), we explored the frequency and the degree of the RGR data. To be precise, we converted the data series from its original time domain (with equal daily readings) to the frequency domain using a fast Fourier transform algorithm (FFT)^56^. After the conversion, we separated the frequencies (x-axis, cycles per day, i.e. the frequency of growth) and the magnitude spectrum (y-axis, normalised amplitudes, i.e. the degree of growth) and generated underlying growth patterns of all the monitored lines (**Supplementary Note 8**). Noticeably, for *Paragon* WT, although temperatures and developmental profiles were significantly different between 2015 and 2016, the underlying growth patterns for *Paragon* WT in both years were very similar (**Fig. 6b**). We identified two distinct growth peaks: (1) around 15 days (15.3 days and 15.6 days respectively) and (2) seven-eight days (8.4 days and 7.8 days), indicating that *Paragon* WT is likely to control its underlying growth pattern based on the number of elapsed days instead of other factors such as temperatures.

For Late-DTEM and Early-DTEM NILs whose genetic backgrounds only differ by carrying alleles such as *Ppd-D1a* and *Ppd-1* lof, their growth patterns also contain two peaks (**Fig. 6c**): (1) similar to *Paragon* WT, seven-eight days (7.1 days and 7.7 days) and (2) 23.0 days for Early-DTEM and 15.3 days for Later-DTEM. For other *Paragon* NILs (e.g. Stay-Green mutant and Short-Stem), although the patterns were slightly different, we found that at least one growth peak was close to the region of seven-eight days (**Supplementary Note 8** and **Supplementary Fig. 9**). To verify the FFT approach, we created a hypothetical *Paragon* growth data by combining all *Paragon* NILs across two years as a technical replicate. Figure 6d shows that no clear growth peaks can be detected from the hypothetical datasets. Additionally, we have applied the FFT approach to converted the RGR series with equal degree-day readings; similarly, no clear growth peaks can be identified (**Supplementary Note 9**).

As all the tested *Paragon* NILs show a growth peak at seven-eight days and only Late-DTEM had a growth peak at 23 days, this might provide some insights into the mechanism of *Ppd-D1a.* The cyclical 23-day peak in growth over the common 15-day peak might reflect a changed output of the circadian clock of which *Ppd1* (PRR7 in *Arabidopsis*^57^) is accelerating the development in a cyclical manner. We are currently conducting a number of experiments, from gene expression to cell biology to advance our understanding of this discovery generated by the CQ platform.

### Use case 3 – GxE predictive modelling

Crop modelling is used in breeding and crop research for integrating complex external and internal variables to understand GxE interactions and genetic systems. Many existing models use genotypes **(G)** and environmental factors **(E)** as input parameters to predict phenotypes **(P)** as the output of the models^58–62^. Similarly, we established a light-weight GxE model to predict crop growth using continuous crop-climate data collected by the CQ platform. Also, we used the computational power of CQ’s single-board computers to explore how to operate the GxE model on a daily basis together with infield phenotyping tasks.

The key input components (environmental factors, growth stages and growth traits) of the model and how it was utilised for predicting growth in fluctuating growing conditions are summarised in **Figure 7**. Firstly, we selected environmental factors that were strongly correlated with the performance-related traits such as RGR and height at four key growth stages (from jointing to flowering) using *Pearson* correlation (**Supplementary Table 5**). This approach has identified five out of the 12 factors (p<0.01), including degree-day, solar radiation, rainfall, temperature, and daily light duration. Two heat maps (**Fig. 7a**) were produced to present the selected factors at the four stages (Online Methods and **Supplementary Note 10**). After that, we built a stage-based predictive model using training datasets of growth stages in 2015 and 2015 (Online Methods and **Supplementary Note 11**). We employed support vector machines (SVM)^63^, a popular machine learning algorithm for classification, with radial basis function kernels to classify growth stages. **Figure 7b** illustrates the classified growth stages (coloured blue) benchmarked against the stages scored by expert crop physiologists (coloured red). **Supplementary Figure 10** illustrates the performance of the model when classifying the timing and duration for other wheat genotypes. We found that SVMs trained on two-season *Paragon* WT data had the highest scores using the benchmarking approach. Although the model modestly mistimes in booting (GS41-49) and heading (GS51-59) due to their short duration, we are adding new training data acquired in 2016 and 2017 to improve the model.

On the basis of the identified environmental factors and growth stage modelling, we explored a set of linear regression models to establish a global predictive model to forecast the continuous growth curve of *Paragon* genotypes, an approach that can be used to help farmers and breeders to optimise crop growth and genotype selection in the future. Figure 7c shows how the growth predictive model performs on the hypothetical *Paragon* growth data (a technical replicate with mean squared error: 46.593, correlation: 0.999). The environmental factors and corresponding daily RGR data are grouped together for each stage and a linear regression model is then constructed to fit growth rate within the forecast growth stages, together with an ordinary least squares method to determine model coefficients. A relative growth rate estimate equation 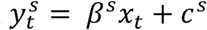 is used, where *x_t_* is the environmental data at time point *t, ß^s^* is the model coefficients (weight vectors), and *c^s^* are constant offsets (i.e. intercepts for each growth stage). The super-script *s* denotes different growth stages (Online Methods and **Supplementary Note 12**).

Growth estimates at each stage are concatenated to form a single vector, based on which height values (in centimetres) and stage-based growth rates are calculated. Using the predictive model, we produced growth estimates for four wheat NILs (**Supplementary Fig. 11**). In this way, we compared how well the model performed with respect to the trait analyses recordings obtained from the CQ platform. To link the predictive model with crop agronomy, we also calculated the average standard deviation (SD) of the predicted crop height values, so a real-time warning message can be triggered on the CropMonitor control system. For example, if crop is growing outside the safe region (the bounds of its estimated height region, ±1SD, **Fig. 7d**), warning messages will be generated to inform the users that the growth is either too quick or too slow.

## Discussion

With the development of modern high-throughput and low-cost genotyping platforms, the current bottleneck in breeding and crop research lies in phenotyping. Here, we describe CropQuant, an automated and scalable field phenotyping system which we believe can enable researchers with a toolkit that can fulfil multi-scale and diverse phenotyping needs for a broader plant research and breeding communities. To deliver the technology, we used the IoT in agriculture ethos to combine networked sensors, distributed computing hardware, computer vision, image analysis, machine learning and modelling to provide high-frequency field phenotyping together with GxE growth predictions, in near real time and in a manner which closely match human scoring.

To enable the CQ system to monitor and compare developmental changes between wheat genotypes using phenotypic data collected under different climate patterns, we carried out an open hardware R&D strategy so that CQ devices could be built and reproduced by other research groups in a relatively easy and cost-effective way. We utilised widely available *Pi* single-board computers and off-the-shelf climate sensors to facilitate the hardware design. For example, a CQ device can be equipped with a range of imaging sensors (e.g. RGB or NoIR *Pi* cameras, or USB and IP cameras) for varied experiments, including using *Pi* cameras to perform side-view and top-view imaging for crop growth and canopy development studies, setting up USB endoscope cameras below the canopy to study tiller abortion, and connecting IP cameras for field level monitoring. Environment sensors were grouped by functions, i.e. ambient and soil-based. So, dissimilar sensor groups could be selected for different experiments. To increase the capability and usefulness of the low-cost CQ device, we provided the hardware construction manual and the circuit board design (**Supplementary Fig. 12**) so that imaging and remote sensing functions could be integrated as well as expanded. We believe that, following the current hardware design, crop-climate data collection in the field can be standardised, data evaluation and communication can be carried in the field, and terminal nodes can be scalable on the CQ platform. Notably, due to the limited manufacturing ability and R&D funding, the maximum indoor and outdoor terminal nodes operating on the CQ platform at the same time was 21, although in theory the system can operate at least 255 nodes simultaneously.

From the software development perspective, we created the CropMonitor control system to provide users with a unified web-based platform which can be used to connect phenotyping hardware with ongoing experiments in an integrated design. Not only does it allow different users to monitor experiments in real time, but it can incorporate different solutions in one shared web place to support experiments at different phases, i.e. from device deployment to the completion of the experiment. With crop-climate data collected in a standard manner, we developed a number of open-source trait analytic algorithms to measure multiple performance-related traits to identify genetic variation under different climate patterns. The software solutions have been evaluated by noisy images caused by complex field environment. Still, it can reliably execute the trait analysis tasks. To verify the results generated by the analysis pipeline, we have scored the performance phenotypes manually on the same plots over two growing seasons and obtained a strong correlation. Furthermore, we established dynamic predictive growth models to forecast the performance of wheat genotypes under varied growing conditions, which could be valuable for agronomic practices. Although the results are promising, it is noticeable that more training datasets are required to improve these models, ideally from varied growing conditions. As the software solutions were implemented on open image analysis, computer vision and machine learning libraries, they can be easily adopted and expanded for other experiments by the communities. To support computational users to understand our work, we have provided detailed comments in our source code.

From a biological perspective, the use of key performance traits generated by the CQ platform can be an excellent tool for screening early establishment, vegetation period, flowering, growth patterns and lodging. For example, vegetative greenness is a useful marker to quantify senescence; utilising the side-view movie, we can closely monitor the process of wheat aging, from the lower stem to the canopy region, a new approach to determine physiological maturity which is important for researching grain development and ripening. Also, continuously monitored greenness can be used in plant pathogen interaction to analyse the activity of pathogens on the leaf surface, as broad yellowish symptoms can be observed from susceptible plants (e.g. rust in wheat). Moreover, crop-climate data acquired by the CQ platform can also assist us to carry out novel biological discoveries. For instance, we are using deep-learning neural network architectures to train a convolutional neural network classifier (CNN) to quantify yield component traits such as spike per unit area and spike/spikelet number (**Supplementary Fig. 13**).

The CQ platform, in combination with networked remote sensors, the web-based control system, computational analytic solutions, and machine-learning based growth modelling, has enabled a cost-effective and scalable field-scale phenotyping of wheat germplasm. Multiple performance-related measurements were quantified in near real time and related to growing conditions. This technology has the potential for multiple applications in breeding and crop production, for example, to optimise the timing of fertiliser applications, irrigation, and predicate harvest dates for maximising yields in different agronomic scenarios. In crop breeding systems, regular field monitoring using the CQ platform identified multiple growth and developmental variables that provided statistically significant phenotypic analysis. These can increase the accuracy of breeding values, particularly for environmental response factors. With more field experimental data collected from different environments feeding into the system, the GxE predictive model and analytics software pipeline can be continuously improved.

In particular, as the field of machine learning has progressed enormously in the last few years, our ability to model complex nonlinear functions and extract high-level phenotypic features is also growing. For example, we are applying deep learning (i.e. CNN and recurrent neural networks, RNNs) to learn and extract features from multidimensional imaging data (including visible and invisible spectrums) that are exceptionally difficult to accomplish through traditional image analysis approaches. Hence, we are consistently exploring deep learning to provide more accurate crop growth and development scores as well as yield-related trait quantifications, offering considerable value to the communities. Our future plan for the CQ platform is to improve the hardware to enhance mobility and modularity, and work with a broader plant research communities to jointly increase the software package for capability and applicability in different growing conditions. So, we could finally deliver real-time infield analysis and integrate field-based phenotyping, UAVs, and satellite into a multi-level and multi-dimensional crop analytic system.

## Conclusion

We believe that the CropQuant technologies described here may have a significant impact on future crop research, breeding activities, and agronomic practices. The reasons are: (1) the low-cost and widely available hardware centred by single-board computers is capable of enabling tasks such as continuous crop monitoring, infield evaluation and efficient data transfer, which are essential for the next-generation field phenotyping; (2) automated trait analysis algorithms integrated in the CQ platform are open-source and expandable software solutions, which are easily accessible and based on community driven numeric and scientific libraries; (3) use cases presented in the paper explain how to apply the CQ platform to study dynamic interactions between genotypes, phenotypes, and environmental factors, which is capable of producing new biological insights of growth patterns through phenotypic analyses. Moreover, our work endeavours to address the affordability and scalability issue for the research communities, which are independent from specific commercial hardware platforms and proprietary or specialised software applications, allowing the utilisation of the CQ platform to accomplish data annotation, performance-related phenotypic analysis, and cross-referencing results freely by the academic communities. Our work confirms previously reported results in the literature and produces novel approaches to enhance the reproducibility of indoor and outdoor crop growth and development experiments. Our case studies of wheat NILs are not limited. Natural variation, mineral or nutrient stress and other crop species could also be monitored using the platform.

## Methods

Methods and any associated references are available in the online version of the paper. Note: Supplementary information is available in the online version of the paper.

## Acknowledgements

We thank members of the Zhou laboratory and the Griffiths laboratory for fruitful discussions. Prof C. Martin and Prof R. Morris at John Innes Centre for reading and improving the manuscript. We thank S. Cossey and Prof Caccamo for supporting the initiation of the project, M. Hewitt for helping to engineer the first prototype, C. Mumford and her team for field trials, NBI infrastructure teams for infield network testing, and research leaders at EI, JIC, TSL and UEA for constructive discussions. This work was strategically funded by the BBSRC, Institute Strategic Programme Grants (BB/J004669/1) and Core Strategic Programme Grant (BB/CSP17270/1) at the Earlham Institute, GRO (BB/J004588/1) to M.W.B. at JIC. J.Z. was partially funded by BBSRC’s Designing Future Wheat Cross-institute Strategic Programme Grants (BB/P016855/1) to Prof G. Moore. The biological research and related software and hardware R&D led by J.Z. was partially supported by BBSRC’s FoF award (GP105-JZ1-B), an NRP Translational Fund (GP069-JZ1-Q), and the UK government’s Eastern Agri-Tech Initiative funding (GP080-JZ1-M).

## Author contributions

J.Z, D.R., S.L., T.L.C., M.D.C., M.W.B. and S.G. designed research; J.Z., D.R., O.G., C.L. and S.O. performed the research; J.Z., D.R., T.L.C., D.W., and G.F. conducted hardware design and the development of analytics software packages; J.Z., D.R., T.L.C., D.W., O.G., C.L., S.L., M.W.B. and S.G. contributed to analyse data; and J.Z., D.R., T.L.C., D.W., M.D.C., M.W.B and S.G. wrote the paper. All authors have read and approved the final manuscript.

## Open Access

The source code is distributed under the terms of the Creative Commons Attribution 4.0 International License (http://creativecommons.org/licenses/by/4.0/), permitting unrestricted use, distribution, and reproduction in any medium, provided you give appropriate credit to the original authors and the source, provide a link to the Creative Commons license, and indicate if changes were made. Unless otherwise stated The Creative Commons Public Domain Dedication waiver applies to the data and results made available in this paper.

## Source code

Source code is freely available for academic usage, which can be downloaded at https://drive.google.com/drive/folders/0B17ZL8AzLo8wNFJUVS1lOFkzb3M?usp=sharing (an online Github repository is being prepared and will be updated in bioRxiv as soon as possible)

## Online Methods

**Five wheat NILs** used in the field trial represent a range of genetic variation all with the genetic background of the UK elite spring wheat ‘*Paragon*’. The development of the Late-DTEM: Par (Norstar + Gamma 319c) 3c-11, *Ppd-1* loss of function (lof) lines is described previously^64^. The development of the Early-DTEM NILs: Par (GS100 2A+CS2B+Son64 2D)-T10 B10 –3b16 and *Ppd-D1a* photoperiod insensitive has also been published^65^. The novel line Stay-Green is line 2316b selected on the basis of stay green phenotype from a population of 7000 *Paragon* EMS mutants carried through single seed descent up to M6 developed under the Wheat Genetic Improvement Network of the UK Department of Food and Rural Affairs (Defra). The semi-dwarf NILs (short) were produced by marker assisted backcrossing (to BC6) using *Rht-B1* and *Rht-D1* KASP markers (LGC). which is available online from http://www.cerealsdb.uk.net/cerealgenomics/CerealsDB. The sources of *Rht-D1b* and *Rht-B1b* were the UK winter wheat varieties ‘Alchemy’ and ‘Robigus’ respectively. The five wheat lines were sown in single 1 m^2^ plots in autumn 2014 at Church Farm, Norfolk UK, and grown according to standard agronomic practice. The manual score of days to ear emergence (DTEM) was done when 50% of the plot showed 50% emergence of the ear from the flag leaf. The manual measurement of plant height was done from the ear tip to ground level.

**The CropQuant hardware** contains many components, of which the centre one is a *Raspberry Pi* 2 or *Pi* 3 single-board computer (we have also used Intel® Edison in a different version of CropQuant workstation). Based on a mobile ARM processor, the *Raspberry Pi* computer features on-board external connections in the form of USB and Ethernet to allow expansion using additional peripherals as well as an array of digital GPIO (general purpose input and output) pins to interface with. The crop growth image acquisition was performed using a 5MP RGB or NoIR (No InfraRed filter) camera module connected via a CSI (Camera Serial Interface) port on the *Pi* mother board. Digital temperature and humidity sensors are connected via manufacturer supplied circuits to the GPIO pins of the *Pi* for interactive control. The sensors themselves are mounted separate from the circuits, externally on the CropQuant’s housing, wired through the base of the device and sheltered by a smaller, open housing unit. The external mounting allows for accurate sensing of ambient air conditions while sheltering the electronics from direct water damage. The CropQuant terminal is housed within a weatherproof (IP66 rated) plastic container, sealed around all openings allowing operation in the field. Physical connection to the system for data transfer via USB or Ethernet and power (12/5V DC) is facilitated by water-resistant couplers designed to be sealed against the rain and air moisture.

**The CropQuant software package** runs on Linux-based operating system *Debian*. It contains two servers, NetATalk and VNC sever, to facilitate infield data transfer and remote systems control, which allows users to connect to each CQ terminal through a wireless (using a tablet or a smartphone) or a wired connection (using a laptop). To enable real-time systems interactions, a GUI-based imaging program has been developed and added into the software package to control the RGB or NoIR camera module for time-lapse crop monitoring. The program can automatically detect the IP address of a given CropQuant terminal so that the terminal can be associated with its specific experiment ID of the field trial. After that, the program requests users to specify information such as genotype, biological replicates and imaging duration via a GUI dialog box, where users can initiate the image acquisition. The program can automatically adjust white balance, exposure mode and shutter speed in relation to variable infield lighting conditions using the *picamera* package, a Python interface to the *Raspberry Pi* camera hardware. Both image resolution and imaging frequency (three times per hour in our field trials) can be changed if users want to modify their experimental settings. The program also conducts the initial quality control and data backup after each image is captured.

Besides the image acquisition, the software package contains a variety of functions such as performing simple workstation and network diagnostics and synchronising with the central server twice within an hour to upload sensor data and CropQuant hardware information (see CropMonitor). Representative daily images are routinely selected and transferred to the central server during the night, which provides a daily snapshot of the monitored crops. Image data backups held on the SD card of the device are routinely synchronised with the server to provide an external backup as needed, with verification of multiple separate backups being performed by the process before it removes the archived data to free storage space. Relying on the Linux *crontab* scheduling system, we can monitor the performance of the software package and resume it automatically in cases of software interruption or power disruption. The SD card image running on the current version of CropQuant can be downloaded via https://drive.google.com/drive/folders/0B17ZL8AzLo8wNFJUVS1lOFkzb3M?usp=sharing. Source code is freely available for academic usage, which was arranged into source trees and saved in both local and central repositories. We are also preparing an online Github repository for the CropQuant project.https://picamera.readthedocs.io/en/release-1.13/

**CropMonitor** is an IoT-style control system developed to oversee the whole CQ platform. It is operated through an onsite central server, logging updates received from individual clients, i.e. CQ terminals. A Python application on each workstation is running at regular intervals, scheduled by the native *Cron* Linux command line utility. The application queries the terminal to determine workstation status information such as uptime, network addresses and storage usage. Sensor data and more variable system data such as CPU temperature and the usage of processor and memory are sampled at a higher frequency and a median average of the readings is recorded during the half-hourly query. Once the application has collected all necessary data it is encoded into a JSON data object and transmitted over HTTP to the central server which stores the data in an SQL database running on a HPC cluster. CropQuant status is displayed and automatically updated using a web-based interface, determining whether each node is online by the time of the most recent update. The web interface provides information, including the location of each CropQuant terminal in the field (a field map needs to be uploaded to the central server), graphs of collected terminal and sensor data, and facilitates device configuration, SSH and VNC linking to all active nodes. Nodes within the CropMonitor system are categorised into groups and projects as defined by the user, allowing the organisation of workstations and restriction of access to stored data. CropMonitor provides a centralised real-time monitoring system to administer the network of infield workstations and collate collected data for visualisation, batch processing and annotation.

**The image selection algorithm** is designed to perform speedy assessment of large image datasets captured in field trials by comparing images to a number of fixed criteria. The Python-based algorithm can be executed either on a normal computer or a HPC cluster. All images which meet the analysis standards will be collated. Over 200 GB data have been generated by ten offsite CropQuant terminals in the 2015 season during a 95-day period, with 50GB data were actually analysed after the selection procedure. In turn, an image is measured based on its brightness, sharpness, and shadow percentage, allowing all images which perform above a set of thresholds to be retained for further traits analysis. To determine the brightness of an image, the median value of pixel intensity is taken by transforming the image into HSV colour space. If the median intensity value is lower than a set threshold, the image is culled and not used from this point forward. The image clarity is determined by applying a Sobel edge detection^41^ to the image. The detectable edges are calculated and then correlated with sharpness and exposure range of the image. The result of the clarity detection is also compared to a set threshold, which will disqualify images if they are out of focus or unclear with ill-defined edges. The final image test is of the percentage shadow within the visible area. Dark pixels found in an image with an illumination value of below 20% are either too dark for feature extraction or containing too much shadow in monitored plots. Once all rules have been passed, selected images are included in a result folder with a CSV file recording image metadata for further high-throughput image analysis.

**The plot detection algorithm** detects initial reference positions of monitored plots. The algorithm identifies the coordinates of white reference canes (the plot region) and dark height markers on a ranging pole, using an ensemble of colour-based feature selection on the basis of HSV (hue, saturation and value) and *Lab* non-linear colour space. It also classifies pixels into different groups, including sky, soil between plots, crop canopy, shadow, and plot regions using simple unsupervised machine-learning techniques such as k-means and spectral clustering. After detecting initial reference objects in the image, the algorithm establishes a pseudo 3D reference system that records the 2D coordinates of the plot area, the canopy region, and height markers through a range of feature selection approaches. The pixel-metric conversion is also computed based on height markers on the ranging pole.

**The CropMeasurer algorithm** employs an adaptive intensity and dynamic gamma equalisation to adjust colour and contrast to minimise colour distortion caused by diverse infield lighting. The algorithm tracks geometric differences between the plot on a given image and the initial position. If different, a geometric transformation method will be applied to recalibrate the image, which removes areas outside the plot area and could generate different sizes of black bars to the top of the given image. Within a plot, CropMeasurer tracks the crop height by detecting the visible part of the ranging pole and defines the canopy region through a combined adaptive thresholding and local Otsu threshold methods. Finally, the algorithm applies Harris and Shi-Tomasi corner detection methods^47^ to locate corner-featured points within the canopy region. Red pseudo points are generated to represent the tips of erect leaves, reflective surfaces of curving leaves, heads and the corner points on ears. The main orientation of a given plot is quantified based on an optimised Canny edge detection method^49^, which computes the alignment of crop stems.

**Data interpolation and analysis** have been used to handle minor data loss during the field experiments. Four days’ data gap (at the end of May 2015) has been recorded on a number of offsite CropQuant workstations, which was caused by SD card crash due to short-term battery failure. We used cubic spline interpolation method^66^ to fill the small gap in the phenotypic datasets.

**RGR and FFT conversion** the data is recorded on a daily basis, the maximum frequency component visible is every two days (0.5 cycles-per-day) due to the Nyquist-Shannon sampling theorem^67^. The cycles-per-day can be viewed in the same manner as Hertz, which is known as cycles-per-second and indicates a measure of frequency. We represent the frequency in cycles-per-day as the data were recorded at daily intervals.

**The growth stage predictive model** is the basis of the GxExP model. The model is produced to explore how to predict growth stages for different wheat genotypes in relation to real-time performance traits and environment data. It employs support vector machines (SVM), a popular machine learning technique for classification, with radial basis function kernels to classify growth stages. The performance of the model is tested by *Paragon* WT (G1) growth data from 2015 and 2016. For *Paragon* WT (2015), the model is trained with all other 2015 genotype data, whereas for the 2016 *Paragon* WT datasets, the model is trained with all 2015 data, all models utilise K-Fold^2^ cross-validation for prediction. To simulate the real-world situation for the stage prediction, we did *not* allow the model to obtain knowledge of the previous stages. Hence, the model mainly modestly mistimes booting (GS41-49) and heading (GS51-59) due to the short duration of both stages.

The prediction in comparison with the manually recorded growth stages suggests a successful prediction of the timing and duration across all growth stages for both 2015 and 2016 datasets, except for the short transition period during booting (GS41-49), where the duration of booting is two days short. Due to the limited data points for booting across all genotypes used for training, the model cannot differentiate booting from heading sufficiently. For this matter, we are planning to add training datasets from other varieties such as Watkins and Chinese Spring wheat in other field trials. The stage prediction is trained by *Paragon* WT growth data from both 2015 and 2016: (1) to predict the 2015 *Paragon* WT, the model is trained with all other 2015 NIL growth data; (2) whereas the 2016 *Paragon* WT was based on all 2015 growth data. Through this approach, the model can rectify itself using previous years’ training data. After the training phase, the model utilises K-Fold^68^ cross-validation for the growth stage prediction.

**The GxE interaction model** explores the interactions between the recorded crop growth of five wheat genotypes and a number of environmental factors. Correlations are performed for each environmental factor grouped over three days with the recorded growth data. The reason to group environmental factors into nested three-day periods is to remove outliers and smooth the input data. The correlations are determined for the first four growth stage for five genotypes. The analysis is performed on the grouped data as particular stages (e.g. booting and heading) contain few recorded growth data due to the short duration of both stages were present during the growth. To determine the interactions between relative growth rates (RGR) and environmental factors, we used the formula (*e^RGR^*)^-1^ to convert negative correlation values to positive counterparts, as the RGR series is a decreasing sequence in relation to the increasing nature of growth stages.

Based on significant environmental factors, linear regression models^45^ have been explored and a single linear regression model is selected to estimate RGR of five genotypes in relation to given infield environment conditions. Environmental factors with insignificant correlations (where p > 0.01, with respect to the height over the entire time-series) are removed from the analysis as they provide little predictive power. Ordinary least squares are used to derive the model coefficients. The RGR data is normalised to present percentage changes in height between two consecutive days. To predict the canopy height for a given genotype, environment data at each growth stage is input to the global model. To derive the height of the plant over time, successive application *h_t_* =*h_t-l_*(1+*y_t_*) is applied, where *h_t_* is taken from the above equation, *h_t-l_* is the height of the plant at the previous time-point, and *h*_0_ is equal to the initial height.

The performance of the model is verified by estimating the growth of all five NILs, including the overall paragon growth data (GT). The estimation is displayed with respect to the true canopy height datasets. The mean squared error recorded for G2 (*genotype two,* Late-DTEM), G3 (*genotype three,* Early-DTEM) and G4 *(genotype four,* Stay-Green) shows that the estimated height is close to the true growth curves. However, the error is much larger for G1 *(genotype one, Paragon* WT) and G5 *(genotype five* Short). This is due to the majority of crop growth happens during the early stages (GS32-GS59), estimation deviation during these initial stages could affect the overall height results. As the global predictive model might not be sensitive towards specific genotypes, we are still seeking a better approach to incorporate all genotypes with a similar genetic background into the prediction. The stage predictions are used in the linear regression growth model that could be run on a single-board computer such as a *Pi* computer to give accurate quantifications. We have chosen to establish the predictive growth model based on data produced from the CQ platform and hence did not perform cross-validation tests to offer more rigorous evidence of how well the model will generalise to new data. Given larger datasets containing more biological replicates, conducting cross-validation produces more reliable growth models with increased precision. Warning messages will be triggered via the CQ platform, if the crop growth rate has deviated from the bounds of its estimated growth region (±1SD).

**High definition movies** referred in this manuscript can be freely downloaded at https://drive.google.com/drive/folders/0B17ZL8AzLo8wNFJUVS1lQFkzb3M?usp=sharing (source code is freely available for academic usage and we are also preparing an online Github repository for the CropQuant project).

### Code availability

We used a *Jupyter* Notebook (i.e. the iPython Notebook) to present and explain algorithms and software solutions associated with the CQ project. They are freely available for academic use. Software packages running on CQs, high-throughput trait analysis algorithms, and GxE modelling can be downloaded via GoogleDocs for academic usage and an online Github repository is being prepared. https://drive.google.com/drive/folders/0B17ZL8AzLo8wNFJUVS1lOFkzb3M?usp=sharing

---

**Supplementary figure 1.**
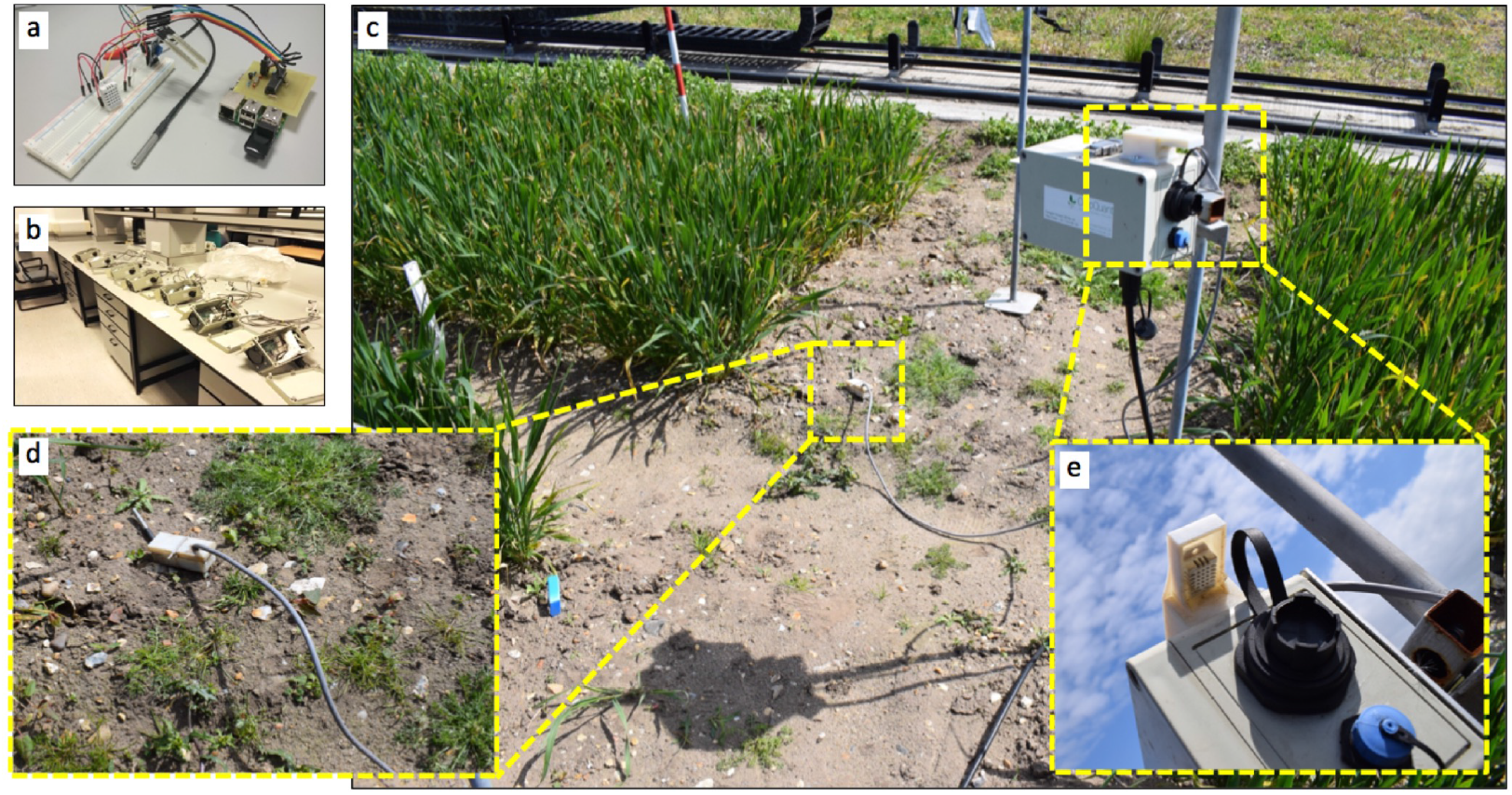
An all-in-one CQ device deployed in field experiments. **(a)** Low-cost remote sensors (e.g. light levels, ambient temperature and humidity, soil temperature and moisture) integrated by a tailored circuit board and then connected to a *Raspberry Pi* computer via GPIO (general purpose input/output) pins. **(b)** A number of all-in-one CQs being tested for establishing a mesh network in the Zhou laboratory. **(c)** An all-in-one CQ device deployed in a wheat field experiment in 2017. **(d)** A soil-based sensor installed to collect soil temperature and moisture for a six-metre wheat plot. **(e)** Light levels and ambient temperature and humidity sensors mounted on the top of the all-in-one CQ, together with an Ethernet coupler (black) and a micro-USB coupler (blue).

**Supplementary figure 2.**
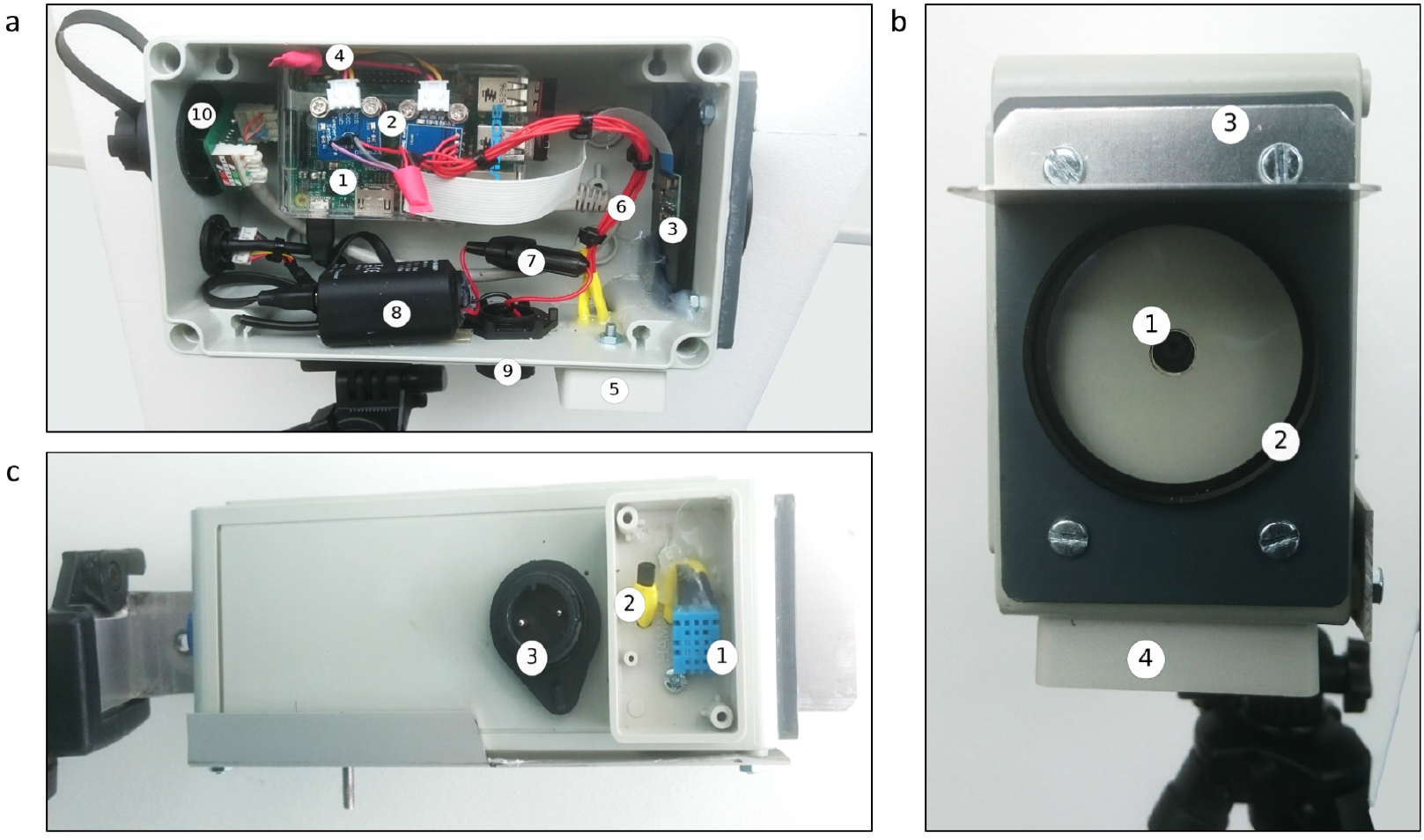
The hardware design of a version 2 CropQuant. **(a)** *Side view:* (1) a *Raspberry Pi* 2, (2) digital sensor circuits, (3) a *Pi* camera module, (4) a sensor GPIO connection, (5) an external sensor housing, (6) a digital sensor connection, (7) an inline power fuse, (8) a voltage converter, (9) an external power connection, (10) an external Ethernet connection coupler. **(b)** *Front view:* (1) a camera module, (2) an external camera UV lens, (3) a camera sunlight shield, (4) an external sensor housing. **(c)** *Base view:* (1) a digital humidity sensor, (2) a digital temperature sensor, and (3) an external power connection.

**Supplementary figure 3.**
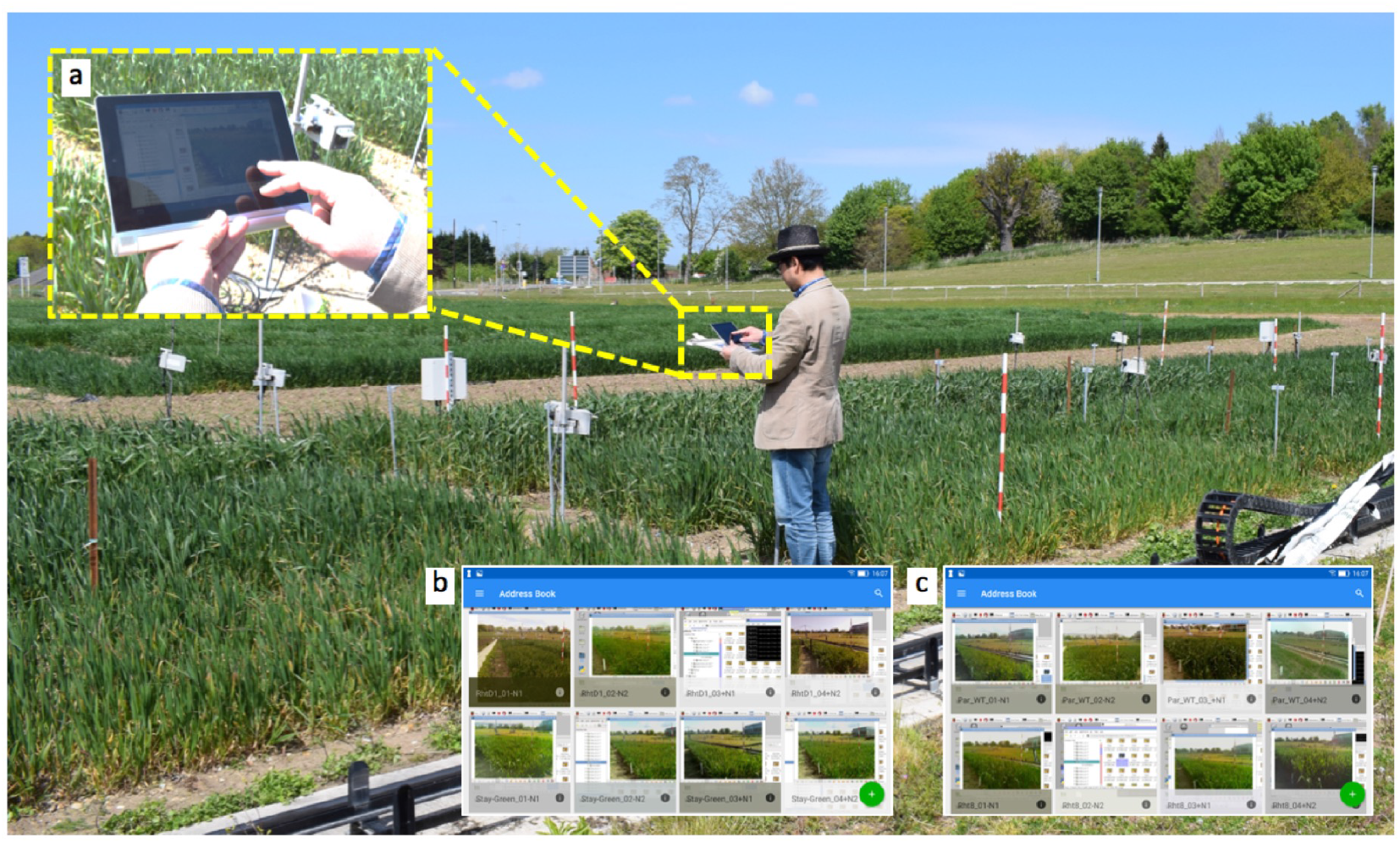
Real-time infield crop monitoring via portable devices. **(a)** A crop scientist using an Android tablet to connect to infield CQ terminals to examine the performance of wheat growth. (b&c) After connecting to the VNC server running on a CQ terminal, different experiments can be inspected and managed by crop scientists via a VNC viewer on the portable device, e.g. smartphones or tablets.

**Supplementary figure 4.**
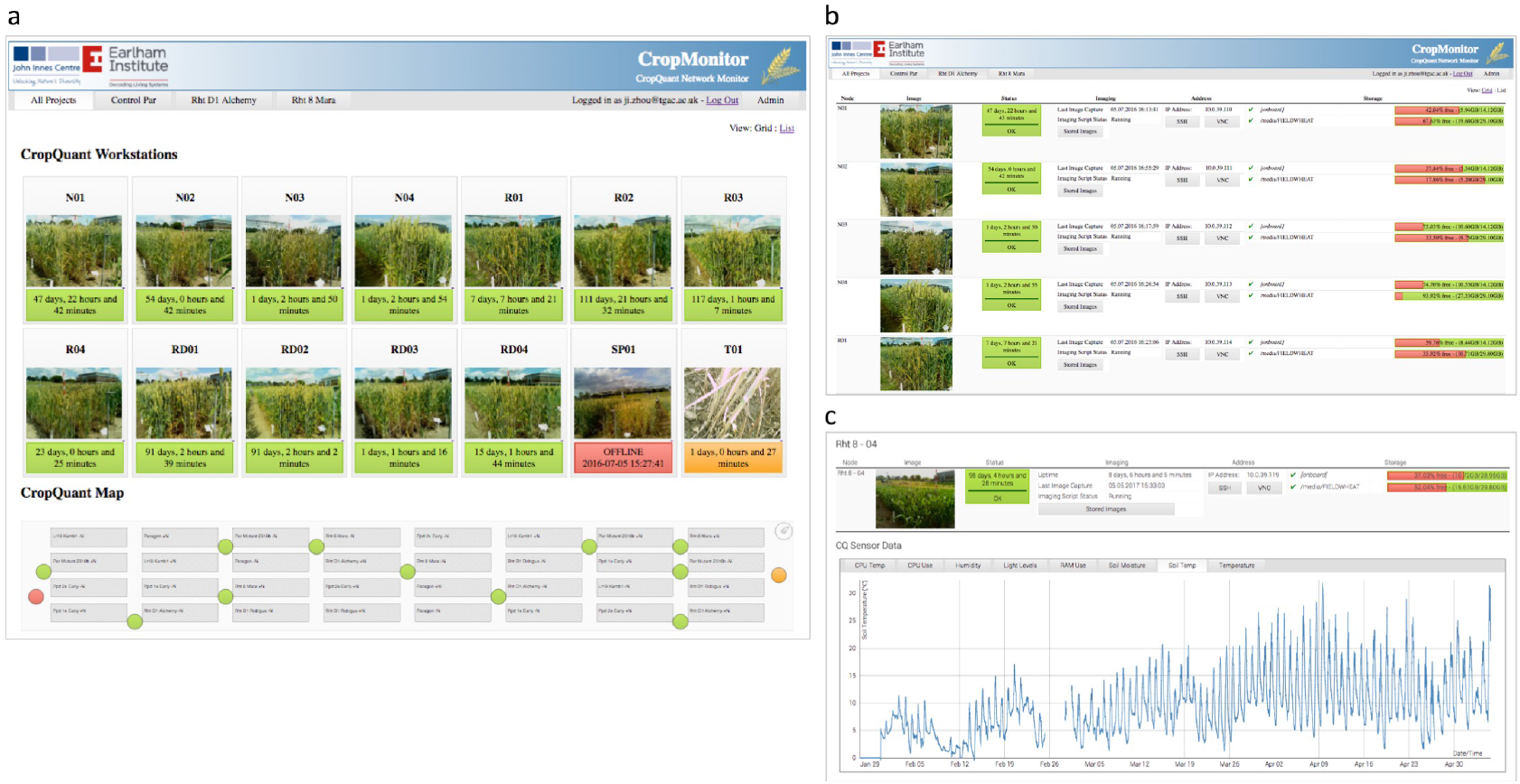
The CropMonitor control system. **(a)** Grid view of the CropMonitor system, which provides a regular overview of all CQ terminals in the field, including online (green) or offline (red) status, operational modes (amber means the imaging is either finished or halted), and the duration of crop monitoring. An experimental layout of monitored plots is also provided, showing the location of all CQ terminals and their operational modes. **(b)** List view of the system showing CQ’s online duration, network addresses, computing storage, and SSH/VNC tools to access CQ terminals directly from the CropMonitor control system. **(c)** Individual view of the system which illustrates an individual CQ workstation, containing climate sensor data and systems information during the monitoring period.

**Supplementary figure 5.**
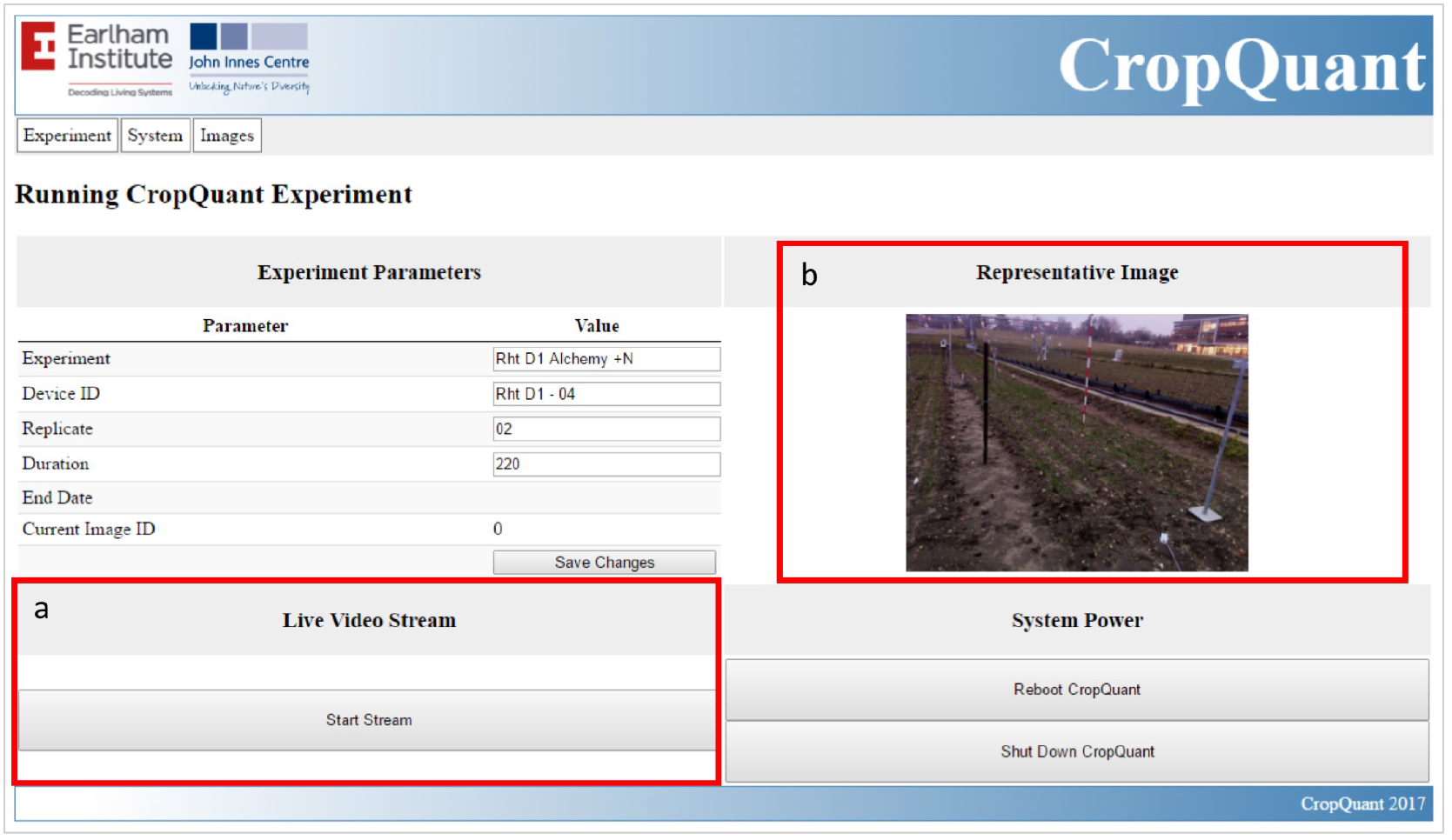
A real-time stream function activated for deploying CQ workstations in the field. (a-b) A live stream function showing the location of a CQ terminal in the field as well as assisting device deployment and systems calibration via the CropMonitor system running on each CQ device.

**Supplementary figure 6.**
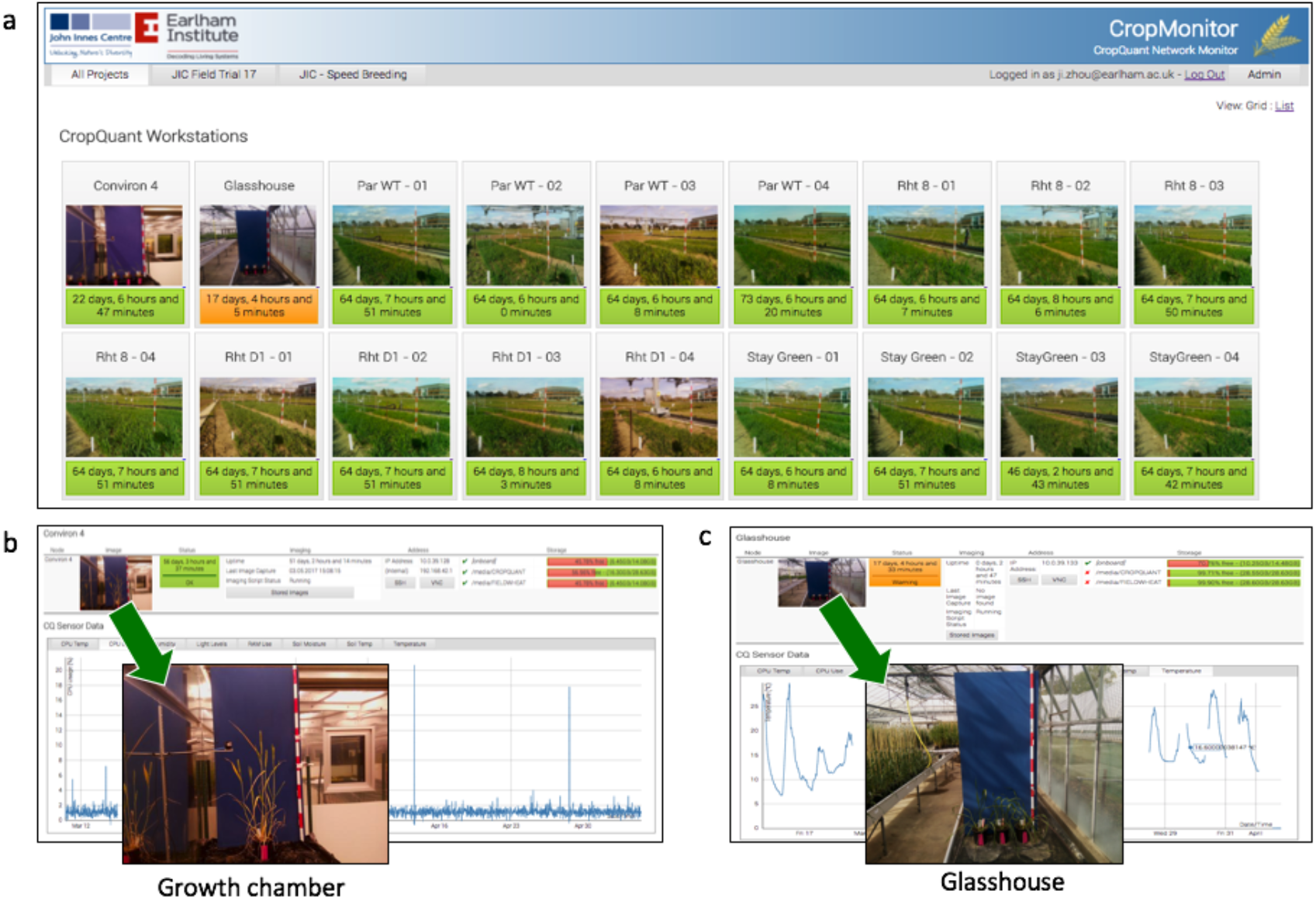
Outdoor and indoor wheat experiments (e.g. wheat speed breeding) monitored by the CQ platform in 2017.

**Supplementary figure 7.**
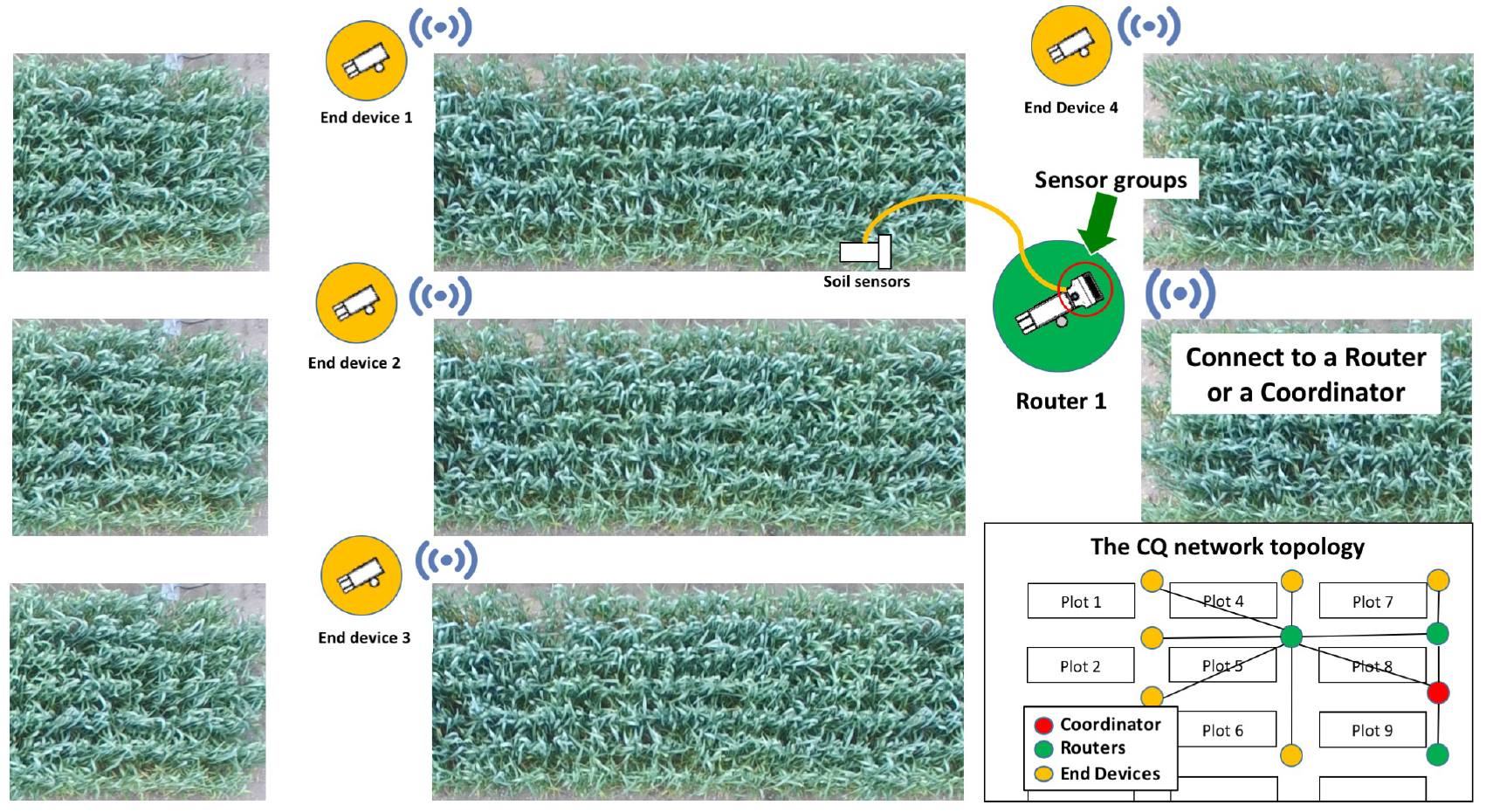
The network topology of the mesh network system established by many all-in-one CQs jointly operating in the field.

**Supplementary figure 8.**
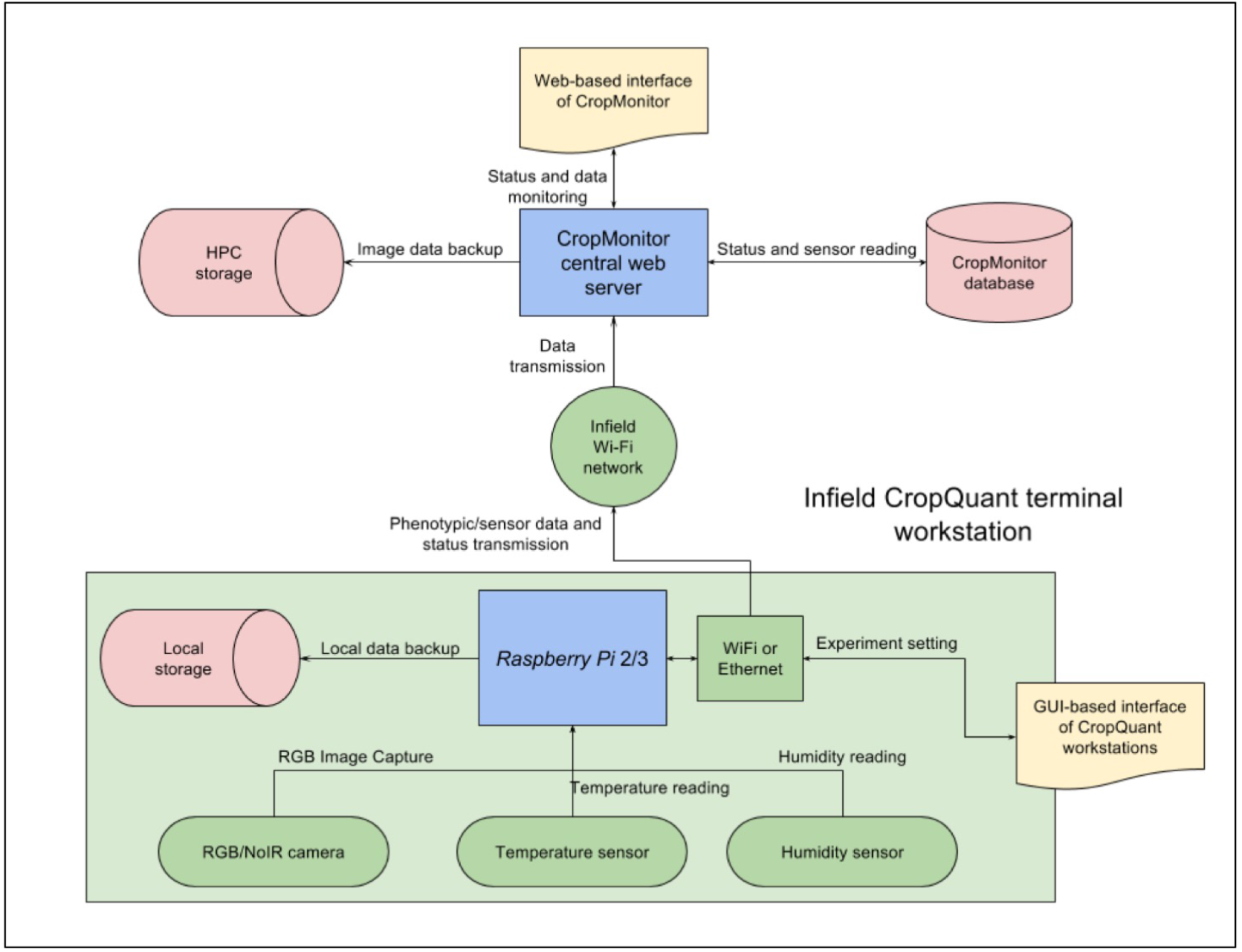
The systems architecture diagram of how to network CropQuant terminals, the CropMonitor system, and a HPC cluster.

**Supplementary figure 9.**
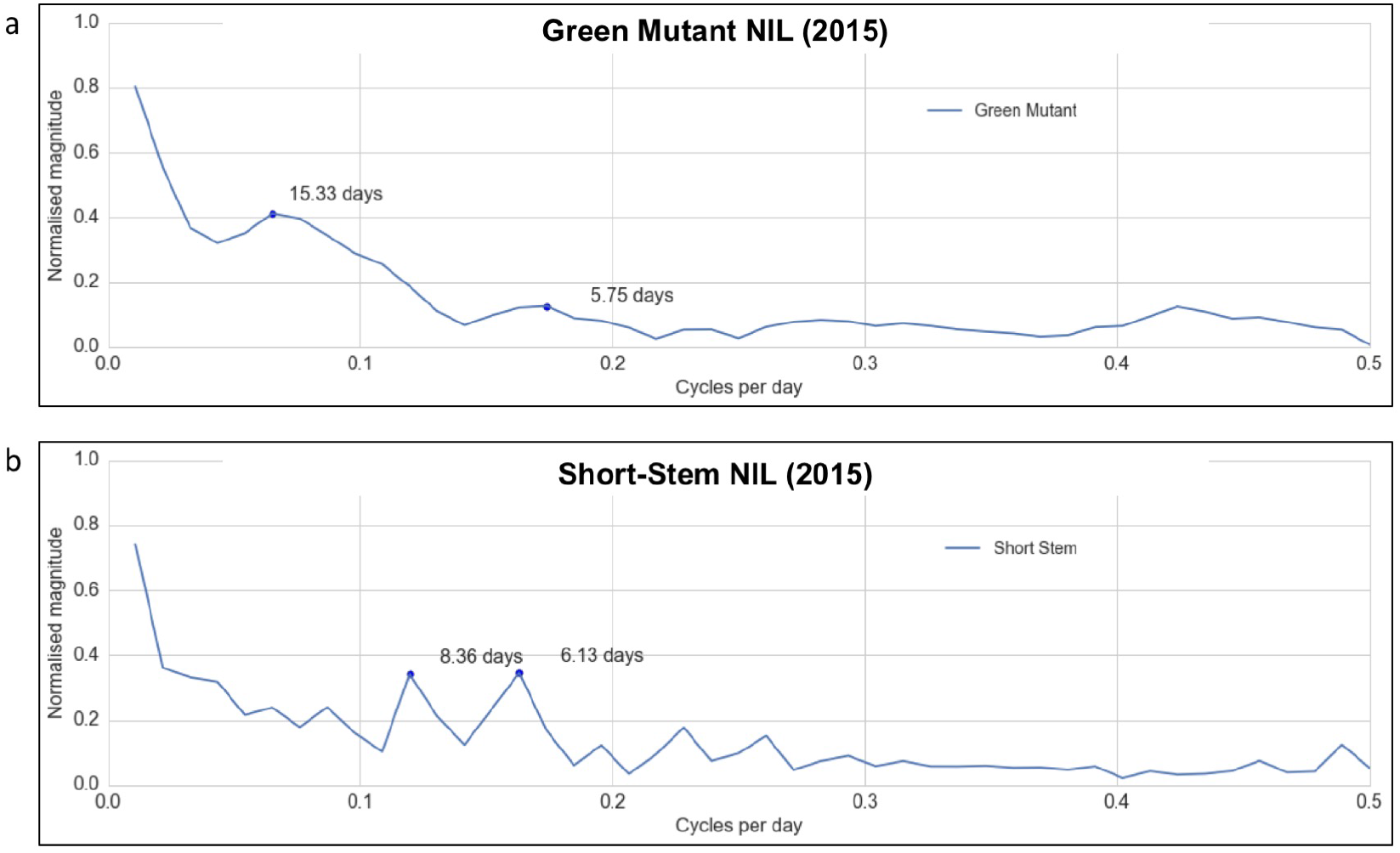
The underlying growth patterns for Green Mutant NIL and Short-Stem NIL extracted through a fast Fourier transform (FFT) approach.

**Supplementary figure 10.**
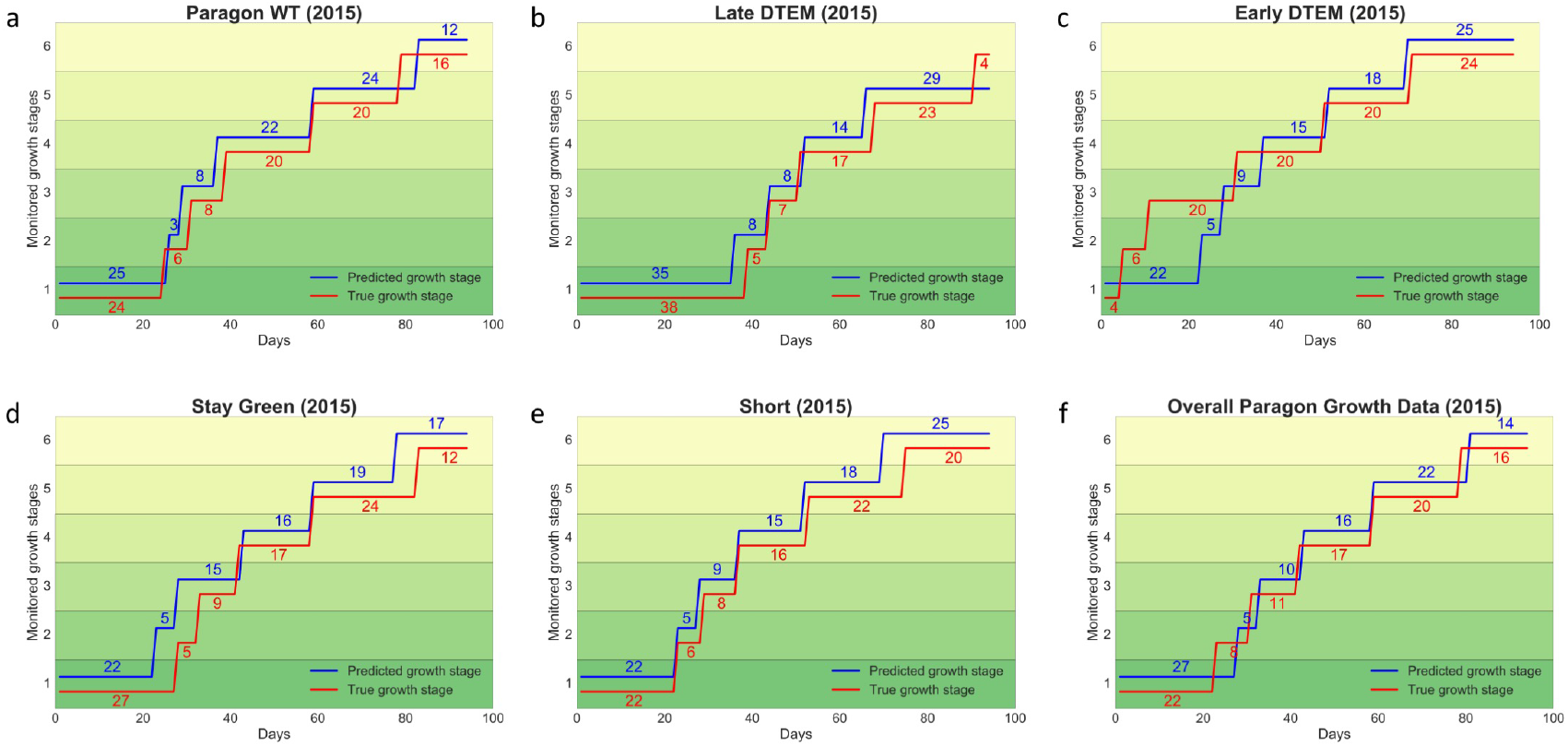
A predictive model to forecast the timing and duration of key growth stages for genotypes with *Paragon* background. (a-e) The model predicts the growth stages based on all *Paragon* growth data (acquired in 2015 and 2016) and environmental factors selected by the GxE correlation model. **(f)** The model forecasts the growth stages based on overall *Paragon* data (the hypothetical growth dataset) and environmental factors.

**Supplementary figure 11.**
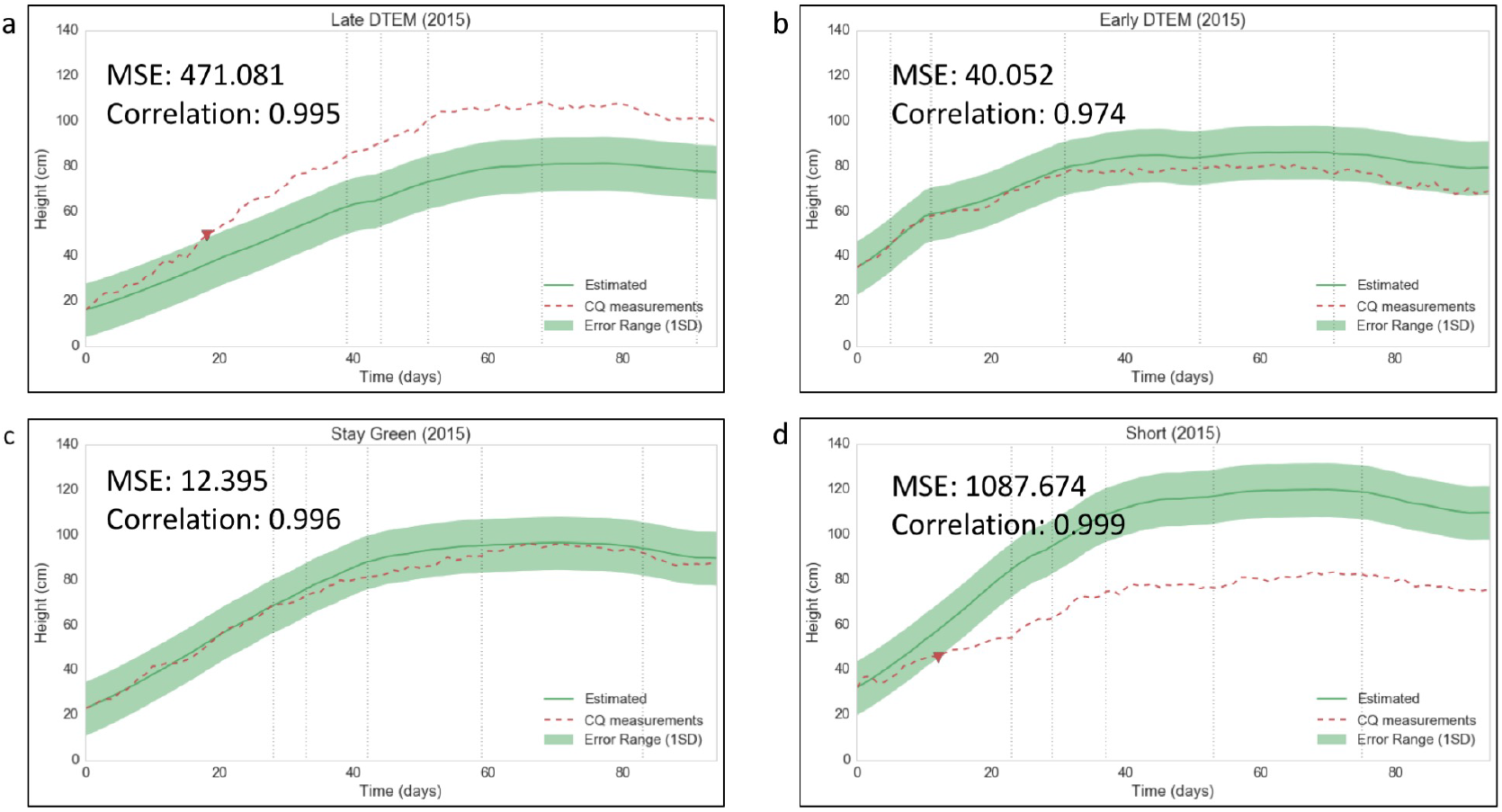
A predictive model to forecast the dynamic growth patterns of wheat genotypes with *Paragon* background in field conditions. (a-b) The growth prediction based on Late-DTEM NIL and Early-DTEM NIL growth data. Vertical dash lines indicate the manually segmented growth stages, from Stem elongation or jointing (GS32-39) to Ripening (GS91-95). The red dotted lines stand for real CQ measurements. If outside the safe bounds of the growth estimates, a warning message (a triangle coloured red) will be triggered on the CropMonitor control system. For example, the Late-DTEM line was growing too slow. **(c)** The growth prediction based on Stay-Green NIL growth data. **(d)** The height prediction based Short-Stem NIL growth data, which was growing too fast.

**Supplementary figure 12.**
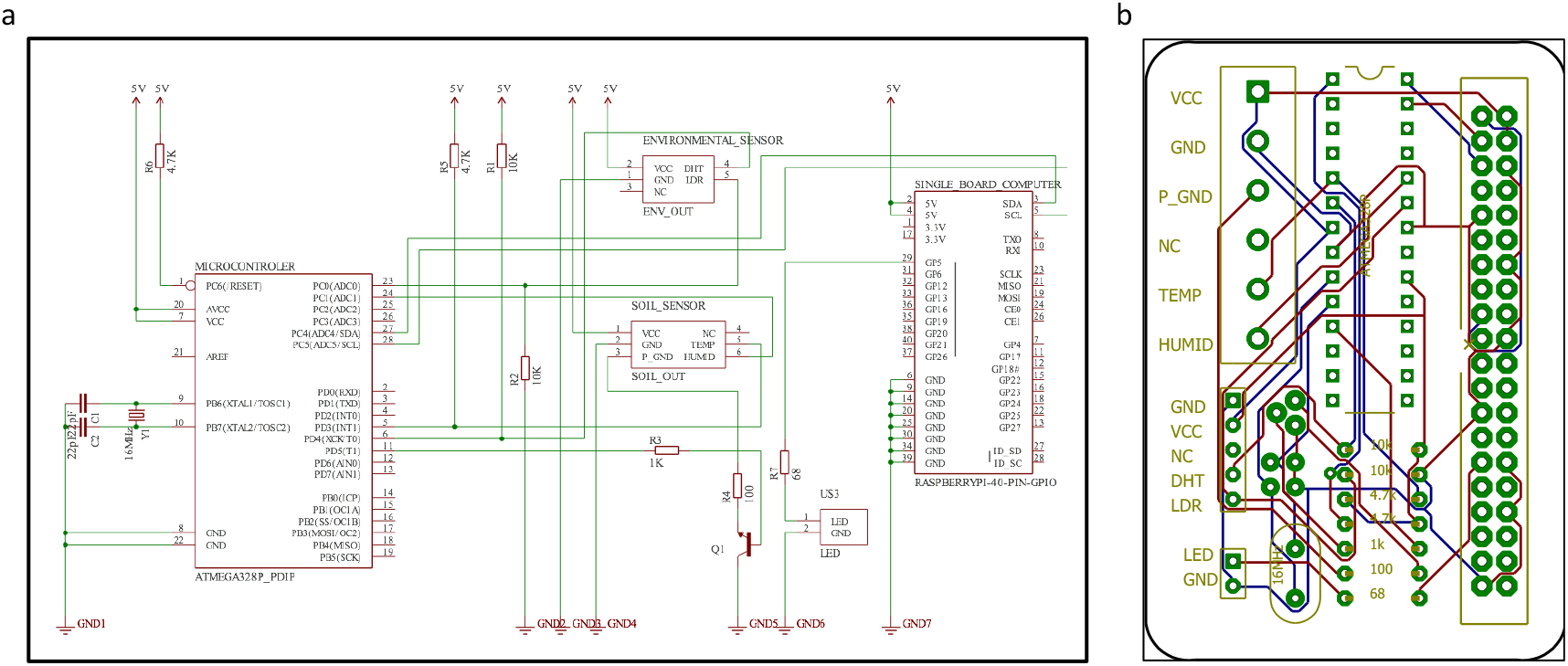
The circuit for the all-in-one CQ workstation.

**Supplementary figure 13.**
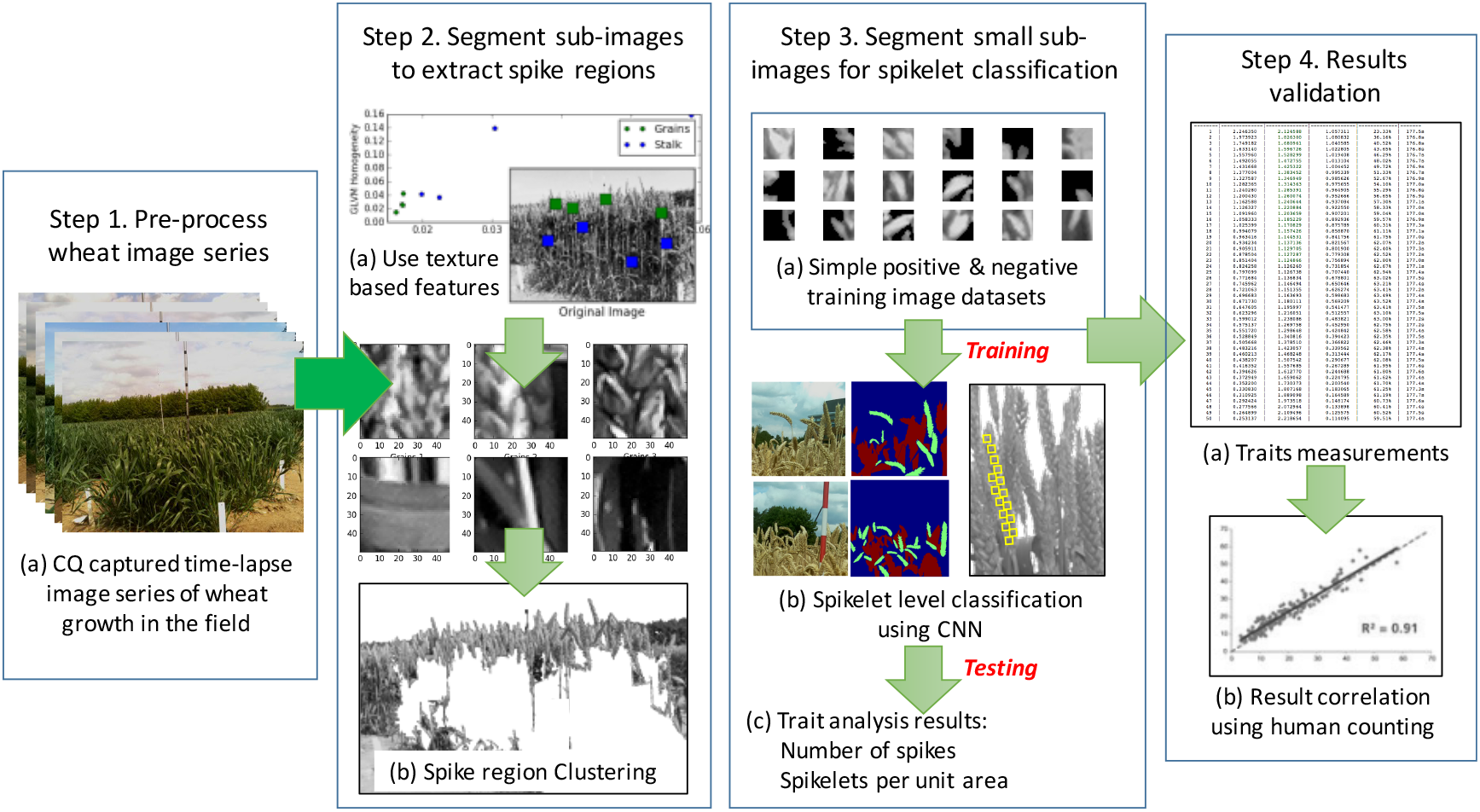
Using deep-learning neural network architectures to train algorithms to quantify yield component traits based on images acquired by the CQ platform. **(1)** *Step 1* pre-processing and calibrate image series captured by the CQ platform in the field. **(2)** *Step 2* based on texture and pattern classification methods such as grey level co-occurrence matrices (GLCM) and texture entropy, detecting spike-featured regions, which are divided into sub-images as training data for a CNN (Convolutional neural network) classifier. **(3)** *Step 3* to count spikes per unit area and spike numbers on a given image, further smaller sub-images are produced to fit a whole spikelet region. CNN is trained to count spikelet-only sub-images. **(4)** *Step 4* the machine-learning based estimation are correlated with manual measurements, so that the estimation model can be improved.

## References

1. Sauer, C. O. Agricultural origins and dispersal. (American Geographical Society, 1952).

2. Donald, C. M. The breeding of crop ideotypes. Euphytica 17, 385–403 (1968).

3. Borlaug, N. E. Wheat breeding and its impact on world food supply. in 3rd International Wheat Genetics Symposium 1–36 (Australian Academy of Science, 1968).

4. Pingali, P. Green Revolution:Impacts, Limits, and the path ahead. Proc. Natl. Acad. Sci. 109, 12302–12308 (2012).

5. Tester, M. & Langridge, P. Breeding Technologies to Increase Crop Production in a Changing World. Science (80-.). 327, 818–822 (2010).

6. Howden, S. M. et al. Adapting agriculture to climate change. Proc. Natl. Acad. Sci. 104, 19691–19696 (2007).

7. McCouch, S. R. et al. Feeding the Future. Nature 499, 23–24 (2013).

8. Malosetti, M., Ribaut, J. M., Vargas, M., Crossa, J. & Van Eeuwijk, F. A. A multi-trait multienvironment QTL mixed model with an application to drought and nitrogen stress trials in maize (Zea mays L.). Euphytica 161, 241–257 (2008).

9. Houle, D., Govindaraju, D. R. & Omholt, S. Phenomics: the next challenge. Nat. Rev. Genet. 11, 855–866 (2010).

10. Hawkesford, M. J. & Lorence, A. Plant phenotyping: increasing throughput and precision at multiple scales. Funct. Plant Biol. (2017). doi:10.1071/FP

11. Barabaschi, D. et al. Next generation breeding. Plant Sci. 242, 3–13 (2015).

12. Yang, W. et al. Combining high-throughput phenotyping and genome-wide association studies to reveal natural genetic variation in rice. Nat. Commun. 5, 1–9 (2014).

13. Brenchley, R. et al. Analysis of the bread wheat genome using whole-genome shotgun sequencing. Nature 491, 705–10 (2012).

14. Araus, J. L. & Cairns, J. E. Field high-throughput phenotyping: the new crop breeding frontier. Trends Plant Sci. 19, 52–61 (2014).

15. Bevan, M. W. et al. Genomic innovation for crop improvement. Nature 543, 346–354 (2017).

16. Furbank, R. T. & Tester, M. Phenomics–technologies to relieve the phenotyping bottleneck. Trends Plant Sci. 16, 635–44 (2011).

17. Cobb, J. N., DeClerck, G., Greenberg, A., Clark, R. & McCouch, S. Next-generation phenotyping: Requirements and strategies for enhancing our understanding of genotype-phenotype relationships and its relevance to crop improvement. Theor. Appl. Genet. 126, 867887 (2013).

18. Deery, D., Jimenez-Berni, J., Jones, H., Sirault, X. & Furbank, R. Proximal Remote Sensing Buggies and Potential Applications for Field-Based Phenotyping. Agronomy 4, 349–379 (2014).

19. Eliceiri, K. et al. Biological imaging software tools. Nature methods 9, (2012).

20. Sankaran, S. et al. Low-altitude, high-resolution aerial imaging systems for row and field crop phenotyping: A review. Eur. J. Agron. 70, 112–123 (2015).

21. Duan, T. et al. Comparison of ground cover estimates from experiment plots in cotton, sorghum and sugarcane based on images and ortho-mosaics captured by UAV. Funct. Plant Biol. (2016). doi:10.1071/FP16123

22. Yang, C., Everitt, J. H., Du, Q., Luo, B. & Chanussot, J. Using high-resolution airborne and satellite imagery to assess crop growth and yield variability for precision agriculture. Proc. IEEE 101, 582–592 (2013).

23. Zaman-Allah, M. et al. Unmanned aerial platform-based multi-spectral imaging for field phenotyping of maize. Plant Methods 11, 35 (2015).

24. Lobell, D. B. The use of satellite data for crop yield gap analysis. F. Crop. Res. 143, 56–64 (2013).

25. Pask, A., Pietragalla, J. & Mullan, D. Physiological Breeding II: A Field Guide to Wheat Phenotyping. CIMMYT (CIMMYT, 2012). doi:10.1017/CBO9781107415324.004

26. Barker, J. et al. Development of a field-based high-throughput mobile phenotyping platform. Comput. Electron. Agric. 122, 74–85 (2016).

27. Andrade-Sanchez, P., Gore, M. & Heun, J. Development and evaluation of a field-based high-throughput phenotyping platform. Funct. Plant Biol. 41, 68–79 (2014).

28. Vadez, V. et al. LeasyScan: A novel concept combining 3D imaging and lysimetry for high-throughput phenotyping of traits controlling plant water budget. J. Exp. Bot. 66, 5581–5593 (2015).

29. Virlet, N., Sabermanesh, P., Sadeghitehran, K. & Hawkesford, M. J. Field Scanalyzer: An automated robotic field phenotyping platform for detailed crop monitoring. Funct. Plant Biol. 44, 143–153 (2016).

30. Chen, D. et al. Dissecting the Phenotypic Components of Crop Plant Growth and Drought Responses Based on High-Throughput Image Analysis. Plant Cell Online 26, 4636–4655 (2014).

31. White, J. W. et al. Field-based phenomics for plant genetics research. F. Crop. Res. 133, 101112 (2012).

32. Kelly, D. et al. An opinion on imaging challenges in phenotyping field crops. Mach. Vis. Appl. 27, 681–694 (2016).

33. Ribaut, J.-M., de Vicente, M. C. & Delannay, X. Molecular breeding in developing countries: challenges and perspectives. Curr. Opin. Plant Biol. 13, 213–218 (2010).

34. Moose, S. P. & Mumm, R. H. Molecular plant breeding as the foundation for 21st century crop improvement. Plant Physiol. 147, 969–977 (2008).

35. Gubbi, J., Buyya, R., Marusic, S. & Palaniswami, M. Internet of Things (IoT): A Vision, Architectural Elements, and Future Directions. Futur. Gener. Comput. Syst. 29, 1645–1660 (2013).

36. the UK Government Chief Scientific Adviser. The Internet of Things: making the most of the Second Digital Revolution. (2014). doi:GS/14/1230

37. Watson, A. et al. Speed breeding: a powerful tool to accelerate crop research and breeding. 117 (2017). doi:dx.doi.org/10.1101/161182

38. Howse, J. OpenCVComputer Vision with Python. (Packt Publishing Ltd., 2013).

39. Pedregosa, F. et al. Scikit-learn: Machine Learning in Python. J. Mach. Learn. Res. 12, 28252830 (2011).

40. van der Walt, S. et al. Scikit-image: image processing in Python. PeerJ 2, 1–18 (2014).

41. Szeliski, R. Computer Vision: Algorithms and Applications. (Springer Science & Business Media, 2010). doi:10.1007/978-1-84882-935-0

42. Aloise, D., Deshpande, A., Hansen, P. & Popat, P. NP-hardness of Euclidean sum-of-squares clustering. Mach. Learn. 75, 245–248 (2009).

43. Shi, J. & Malik, J. Normalized cuts and image segmentation. IEEE Trans. Pattern Anal. Mach. Intell. 22, 888–905 (2000).

44. Bassiou, N. & Kotropoulos, C. Color image histogram equalization by absolute discounting back-off. Comput. Vis. Image Underst. 107, 108–122 (2007).

45. Sezgin, M. & Sankur, B. Survey over image thresholding techniques and quantitative performance evaluation. J. Electron. Imaging 13, 146–165 (2004).

46. Haralick, R., Shanmugan, K. & Dinstein, I. Textural features for image classification. IEEE Transactions on Systems, Man and Cybernetics 3, 610–621 (1973).

47. Harris, C. & Stephens, M. A Combined Corner and Edge Detector. Procedings Alvey Vis. Conf. 1988 147–151 (1988). doi:10.5244/C.2.23

48. Zadocks, J. C., Chang, T. T. & Konzak, C. F. A decimal code for the growth stages of cereals. Weed Res. 14, 415–421 (1974).

49. Kimmel, R. & Bruckstein, A. M. Regularized Laplacian zero crossings as optimal edge integrators. Int. J. Comput. Vis. 53, 225–243 (2003).

50. Semenov, M. A. & Doblas-Reyes, F. J. Utility of dynamical seasonal forecasts in predicting crop yield. Clim. Res. 34, 71–81 (2007).

51. Griffiths, S. et al. Meta-QTL analysis of the genetic control of ear emergence in elite European winter wheat germplasm. Theor. Appl. Genet. 119, 383–95 (2009).

52. McMaster, G. S. & Wilhelm, W. W. Growing degree-days: One equation, two interpretations. Agric. For. Meteorol. 87, 291–300 (1997).

53. Lancashire, P. D. et al. A uniform decimal code for growth stages of crops and weeds. Ann. Appl. Biol. 119, 561–601 (1991).

54. Delude, C. M. Deep phenotyping: The details of disease. Nature 527, S14–S15 (2015).

55. Fiorani, F. & Schurr, U. Future scenarios for plant phenotyping. Annu. Rev. Plant Biol. 64, 267–91 (2013).

56. Heideman, M., Johnson, D. & Burrus, C. Gauss and the history of the fast fourier transform. IEEE ASSP Mag. 1, 14–21 (1984).

57. Nakamichi, N., Kita, M., Ito, S., Yamashino, T. & Mizuno, T. PSEUDO-RESPONSE REGULATORS, PRR9, PRR7 and PRR5, Together play essential roles close to the circadian clock of Arabidopsis thaliana. Plant Cell Physiol. 46, 686–698 (2005).

58. Chenu, K., Deihimfard, R. & Chapman, S. C. Large-scale characterization of drought pattern: A continent-wide modelling approach applied to the Australian wheatbelt – spatial and temporal trends. New Phytol. 198, 801–820 (2013).

59. Tardieu, F. Why work and discuss the basic principles of plant modelling 50 years after the first plant models? J. Exp. Bot. 61, 2039–2041 (2010).

60. Turc, O., Bouteill, M., Fuad-Hassan, A., Welcker, C. & Tardieu, F. The growth of vegetative and reproductive structures (leaves and silks) respond similarly to hydraulic cues in maize. New Phytol. 212, 377–388 (2016).

61. Dawson, I. K. et al. Barley: a translational model for adaptation to climate change. New Phytol. 206, 913–931 (2015).

62. Palosuo, T. et al. Simulation of winter wheat yield and its variability in different climates of Europe: A comparison of eight crop growth models. Eur. J. Agron. 35, 103–114 (2011).

63. Hsu, C.-W. & Lin, C.-J. A comparison of methods for multiclass support vector machines. IEEE Trans. Neural Networks 13, 415–425 (2002).

64. Shaw, L. M., Turner, A. S., Herry, L., Griffiths, S. & Laurie, D. A. Mutant alleles of Photoperiod-1 in Wheat (Triticum aestivum L.) that confer a late flowering phenotype in long days. PLoS One 8, (2013).

65. Shaw, L. M., Turner, A. S. & Laurie, D. A. The impact of photoperiod insensitive Ppd-1a mutations on the photoperiod pathway across the three genomes of hexaploid wheat (Triticum aestivum). Plant J. 71, 71–84 (2012).

66. R Development Core Team. R: A Language and Environment for Statistical Computing. 1, (R Foundation for Statistical Computing, 2008).

67. Shannon, C. E. Editorial note on ‘Communication in the presence of noise’. Proc. IEEE 72, 1713–1713 (1984).

68. Kohavi, R. A Study of Cross-Validation and Bootstrap for Accuracy Estimation and Model Selection. Int. Jt. Conf Artif Intell. 14, 1137–1143 (1995).

